# Taxor: Fast and space-efficient taxonomic classification of long reads with hierarchical interleaved XOR filters

**DOI:** 10.1101/2023.07.20.549822

**Authors:** Jens-Uwe Ulrich, Bernhard Y. Renard

## Abstract

Metagenomic long-read sequencing is gaining popularity for various applications, including pathogen detection and microbiome studies. To analyze the large data created in those studies, software tools need to taxonomically classify the sequenced molecules and estimate the relative abundances of organisms in the sequenced sample. Due to the exponential growth of reference genome databases, the current taxonomic classification methods have large computational requirements. This issue motivated us to develop a new data structure for fast and memory-efficient querying of long reads. Here we present Taxor as a new tool for long-read metagenomic classification using a hierarchical interleaved XOR filter data structure for indexing and querying large reference genome sets. Taxor implements several k-mer-based approaches such as syncmers for pseudoalignment to classify reads and an Expectation-Maximization algorithm for metagenomic profiling. Our results show that Taxor outperforms competing shortand long-read tools regarding precision, while having a similar recall. Most notably, Taxor reduces the memory requirements and index size by more than 50% and is among the fastest tools regarding query times. This enables real-time metagenomics analysis with large reference databases on a small laptop in the field. Taxor is available at https://gitlab.com/dacs-hpi/taxor.

## Introduction

Identifying organisms in an environmental or clinical sample is a fundamental task in many metagenomic sequencing projects. This includes the detection of pathogens in samples with a large host background (1), as well as studying the composition of microbial communities composed of bacteria, archaea, viruses, and fungi (2). Over the last years, many tools have been developed that classify short and long sequencing reads by comparing their nucleotide sequences with a predefined set of references (3–5). While each tool uses a different classification strategy, they all try to resolve the species present in the sample and determine their relative abundances (6, 7).

Among the different read classification strategies, alignment-based approaches were the first used for taxonomic profiling. Tools like SLIMM (8), DUDes (9) or PathoScope (10) use the results of common read mappers like Bowtie2 (11) and bin the sequencing reads across the different reference genomes. Although these methods have high accuracy, their computational performance decreases tremendously when using entire public databases such as NCBI RefSeq or GTDB as reference datasets. Thus, high-performance computing clusters are needed to run these tools in a reasonable amount of time and fulfill their memory requirements. In contrast, marker-based approaches such as MetaPhlAn2 (12) and mOTUs2 (13) identify bacterial and archaeal species by their 18S or 16S rRNA genes. However, this approach is infeasible for viruses since they have no universally conserved genes. More recent taxonomic classification strategies rely on machine-learning approaches. Tools such as DeepMicrobes (14) and BERTax (15) show promising results for classifying reads on higher taxonomic levels but perform poorly at genus and species levels. Most state-of-the-art taxonomic profilers, like Kraken2 (4), Ganon (16) and KMCP (17), use k-mer-based methods for read classification. In a first step, these methods count the exact matches of substrings of length k among the different reference sequences in the database and use further statistical analysis to assign reads to references. These profiling tools mainly differ in the indexing of the reference set and/or the k-mer selection method used to calculate the similarity between read and reference sequences.

What all taxonomic classifiers have in common is that they struggle with the ever-increasing amount of reference genomes. Databases such as NCBI RefSeq (18) and GTDB (19) already comprise hundreds of thousands of microbial reference assemblies belonging to 62,000 bacterial species (GTDB Release 207) and 12,000 viral species (RefSeq Release 211) and are constantly increasing. This poses a major computational challenge to the profilers in terms of memory usage, index construction, and query time.

Several approaches for efficient indexing and querying of large collections of reference sequence sets have been developed over the last few years to overcome these issues. The popular Kraken2 (4) classifier uses minimizers and introduces a probabilistic, compact hash table to reduce the size of the index. On the other hand, color-aggregative methods like Bifrost (20) or Mantis (21) use compacted de Brujn graphs or counting quotient filters for indexing and querying k-mers. Those methods have the disadvantage that they index each reference separately using data structures for approximate membership queries, e.g. Bloom filters (22). In contrast, Sequence Bloom Tree (SBT) (23–25) approaches exploit the k-mer redundancy in homogeneous datasets such as those from RNA-Seq experiments to compare large sequence datasets. However, these approaches are unsuitable for heterogeneous k-mer sets, such as microbial genomes. Tools like BIGSI (BItsliced Genomic Signature Index)(26), COBS (Compact Bit-Sliced Signature Index) (27) and KMCP (17), which are based on Bloom filter matrices, are promising much better results for taxonomic profiling of large sequencing data sets. Interleaved Bloom filters (IBF), which belong to the latter approaches, improve the indexing data structures by combining several Bloom filters (one per reference) in an interleaved fashion while allowing to query all Bloom filters at once (28). The IBF data structure has been used by the taxonomic classifier Ganon (16) and was recently enhanced by introducing the Hierarchical Interleaved Bloom Filter (HIBF) in a tool called Raptor (29). Many k-mer-based classifiers use Bloom filters for approximate membership queries of k-mers in large reference data sets. Their popularity is based on their flexibility, low memory requirements, and fast query times. However, there is a small probability that a k-mer is incorrectly reported as being present in a reference sequence, called a false positive. Some tools let the user define the false positive rate and adapt the Bloom filter size and/or the number of hash functions to that value. Although Bloom filters have a low memory footprint, they use 44% more memory than the theoretical lower bound, even when applied in an optimal manner (30). Therefore, several advanced probabilistic filters like cuckoo filters (31, 32) have been developed over the last few years. In particular, XOR filters have been proposed as an alternative to Bloom Filters, using only 23% more memory than the theoretical lower bound (33).

Based on the work of Graf and Lemire and Dadi et al., we first developed an Interleaved XOR filter (IXF) which can be used in the same manner as the Interleaved Bloom Filter implemented as part of the Seqan C++ library (34). We then extended our new data structure to a hierarchical interleaved XOR filter (HIXF) to avoid using more space than necessary when references in the database are highly divergent in size. The HIXF data structure is implemented as part of the taxonomic classification tool Taxor, which allows the user to choose different k-mer-based strategies, such as k-mers and minimizers. Since our new tool is specifically designed for long-read metagenomics experiments, we also implemented open canonical syncmers (OCS) as a k-mer selection approach, which has been shown to be superior to minimizers for error-prone long reads (35, 36). In the final step, taxonomic profiling of the query results is performed by utilizing an expectation maximization (EM) algorithm for abundance estimation and re-assignment of classified reads. We compare Taxor to five state-of-the-art shortand long-read taxonomic classification tools on simulated and real mock communities. Our results show that Taxor can tremendously reduce the index size and memory requirements for queries, while still being on par with the evaluated tools regarding precision and recall.

## Methods

We have designed and implemented our novel taxonomic profiling tool Taxor as a modular workflow that consists of three mandatory steps. First, Taxor computes the k-mer content of the input reference genomes and creates an index for each set of reference genomes. The index is a hierarchical interleaved XOR filter (HIXF), a novel space-efficient data structure for approximate membership queries that we will describe in the following subsections. In the second step, sequencing reads are queried against one or several HIXF index files, resulting in one intermediate file for each index. These intermediate files contain all matches of the reads against the different reference genomes and must be merged before the final profiling step. Three-step filtering of spurious matches is performed before an expectation-maximization algorithm computes taxonomic abundances and reassigns reads based on the taxonomic profile. We finally provide three output files containing information about the sequence abundances (based solely on nucleotide abundance), taxonomic binning (read to reference assignments), and taxonomic abundances (normalized by genome size).

### Interleaved XOR Filter

Many tools for taxonomic classification of sequencing reads facilitate approximate membership queries of k-mers of reads against k-mer sets of the reference sequences. A common approach to implement approximate membership queries is using Bloom filters (22). Dadi et al. improved this approach by developing an Interleaved Bloom Filter that stores several Bloom filters in one single-bit array that allows querying all Bloom filters simultaneously. Inspired by their approach, we developed an Interleaved XOR Filter (IXF) that combines several XOR filters into one data structure, enabling simultaneous querying of all XOR filters. As described by Graf and Lemire, and similar to Bloom filters, the XOR filter uses three independent hash functions that return a corresponding position in the filter for each key (or k-mer). In a Bloom filter, each bit is considered its own array slot, and bits at the positions to which the hash functions point are set to one. In contrast, in an XOR filter, the bits are grouped together into *L*-bit sequences, as shown in Supplementary Figure S1. These *L*-bit sequences in the XOR filter are set in such a way that a bitwise XOR of the three *L*-bit sequences, corresponding to positions returned by the hash functions, equals the result of the *fingerprint* hash function. While building an XOR filter is almost always successful for larger sets of more than |*S* | = 10^7^ elements (37), it can fail for smaller sets, which requires rebuilding with other hash functions. Graf and Lemire have experimentally shown that the estimated probability for the successful building is always greater than 0.8 if the XOR filter size is set to ⌊1.23× |*S* |+ 32⌋, which makes this size constraint an optimal compromise between space requirement and build time.

Our IXF implementation combines several XOR Filters (bins) in one single bitvector, using 8-bit sequences for each XOR filter. Therefore, we first need to initially calculate the size of each XOR filter by ⌊1.23×|*S*| + 32⌋, where *S* is the set of reference k-mers. As for the interleaved Bloom Filter, the largest XOR filter (or largest reference sequence) determines the size of all bins and, thus, also the size of the entire IXF. If *R* is the set of reference sequences to be stored in the IXF and *S*_*r*_ the set of k-mers computed for reference *r*, the size of the IXF can be calculated as follows:

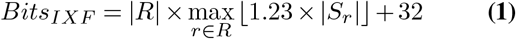

The IXF can be divided into several subvectors, each having the size of the number of bins. Since one bin in the IXF corresponds to exactly one reference sequence, the size of each subvector corresponds to the number of references. When building the Interleaved XOR filter for a set of reference sequences, we compute the k-mer sets for each reference sequence and construct the single XOR filters for each reference according to the algorithm proposed by Graf and Lemire. We used the same hash and fingerprint functions for each XOR filter and combined them in an interleaved fashion, as shown in Fig. 3.

**Fig. 1.**
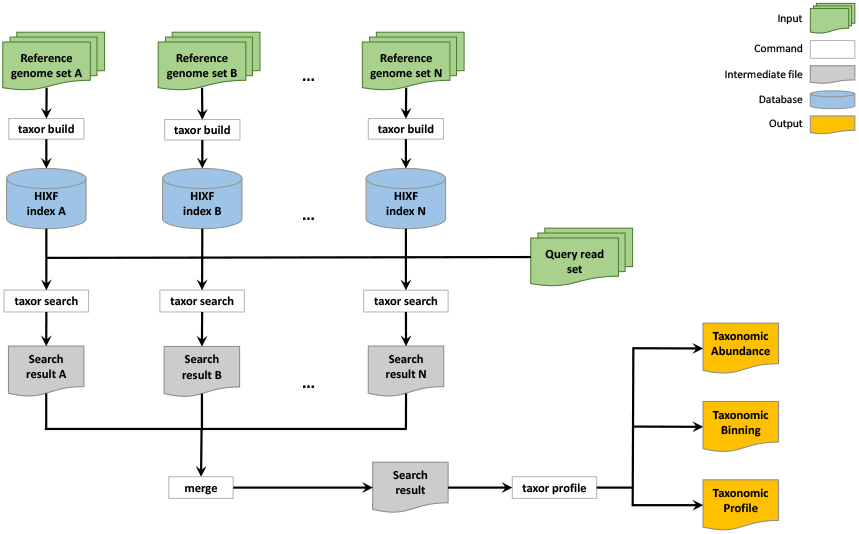
Workflow for taxonomic profiling with Taxor. A typical workflow for using Taxor starts with creating a HIXF index for each reference genome set. Next, a set of sequencing reads is queried against each of the different HIXF index files using the taxor search subcommand, which results in an intermediate search result file for each index file. The search result files must be merged before the subcommand taxor profile calculates the taxonomic profiling result files.

**Fig. 2.**
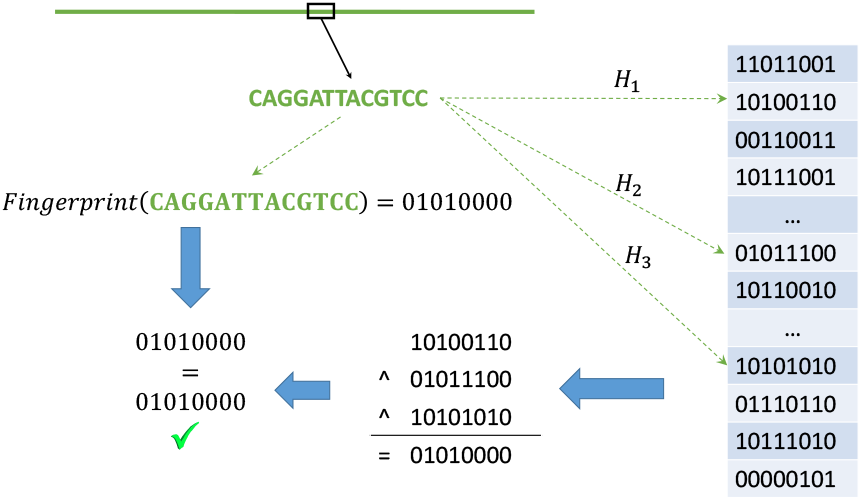
Creating an XOR Filter for a given reference sequence. For each k-mer of the reference sequence an *L*-bit fingerprint and three hash values are calculated. The hash values point to positions in an array consisting of *L*-bit sequences. The *L*bit sequences are set such that a bit-wise XOR of the three *L*-bit sequences equals the result of the fingerprint function for that k-mer. For example, the fingerprint for the 13-mer *CAGGATTACGTCC* equals 01010000, and thus a bitwise XOR of the three 8-bit sequences at positions given by the results of the three hash functions *H*_1_ to *H*_3_ will also result in the 8-bit sequence 01010000

**Fig. 3.**
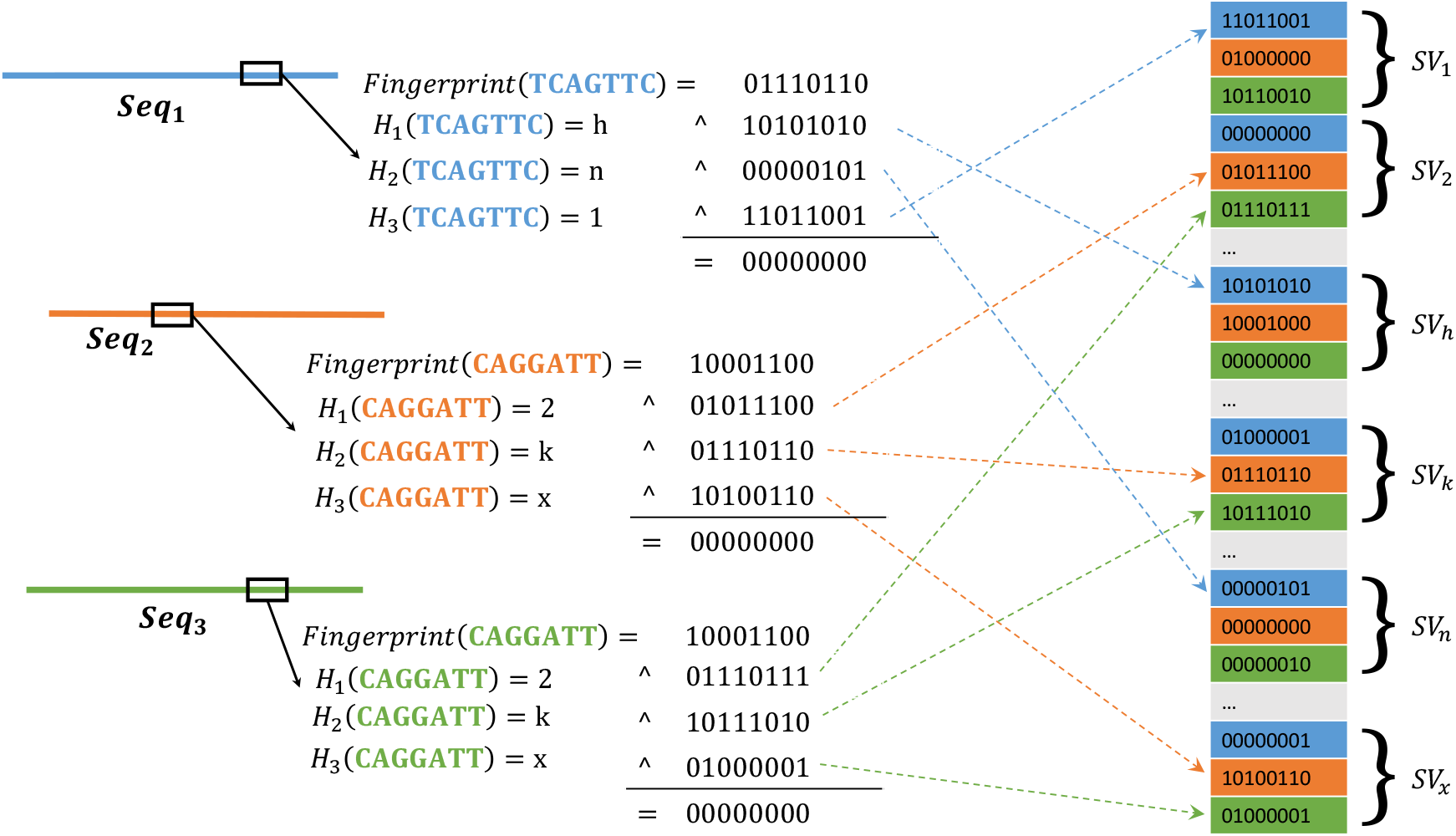
Creating an Interleaved XOR Filter for three reference sequences. For each k-mer of the three differently colored reference sequences, an *L*-bit fingerprint and three hash values are calculated. The hash values determine the three subvectors *SV*_*j*_ in which the *L*-bit sequence of the corresponding position of the reference sequence is set. *L*-bit sequences at those positions are set such that a bit-wise XOR of the three *L*-bit sequences and the fingerprint equal to an *L* − *bit* sequence of zeros. For example, the fingerprint for the 7-mer *TCAGTCC* from *Seq*_1_ equals 01110110, and the three hash functions point to the subvectors *SV*_1_, *SV*_*h*_ and *SV*_*n*_. Now we set the 8-bit sequences at the first position in each subvector such that a bit-wise XOR of the 8-bit sequences and the fingerprint equals to 00000000.

When querying a read against the references stored in the interleaved XOR filter, every k-mer of that read is matched against each XOR filter simultaneously. For each k-mer, we first retrieve the three subvectors *SV*_*j*_ by calculating the three hash values for that k-mer. Next, the resulting *L*-bit sequence calculated by the fingerprint function is concatenated with itself to the length of the subvectors. Applying a logical bitwise XOR to the three subvectors and the fingerprint vector results in a final *L*-bit sequence for each reference. If this sequence equals zero, a bit in a binning bitvector is set to one, indicating the presence of the k-mer in the corresponding reference. Combining the binning bit-vectors of the k-mers to a counting vector finally results in the number of matching k-mers between the read and each reference in the IXF. The example in Fig. 4 visualizes this process. Here, the read consists of two 7-mers, for which we have to calculate the three hash values that point us to the corresponding subvectors *SV*_*j*_, e.g., *SV*_2_, *SV*_*k*_ and *SV*_*x*_ for k-mer CAGGATT. A logical bitwise XOR of the three subvectors and the k-mer’s fingerprint vector results in an 8-bit sequence for each bin. In Fig. 4, the resulting 8-bit sequences of references two (orange) and three (green) equal zero, which yields in setting the bit in the binning bitvector for the references to one. Finally, the counters of the corresponding references are incremented in the counting vector of the read. The resulting counting vector stores the number of matching k-mers between the read and each reference sequence stored in the IXF. Thus, instead of computing three hash values for every XOR Filter separately, we only need to calculate the three hash values once, which poses a significant reduction in computing time to investigate the membership of a k-mer in every XOR Filter. This method lets us quickly count the matching k-mers between the reference genome and a specific sequencing read.

**Fig. 4.**
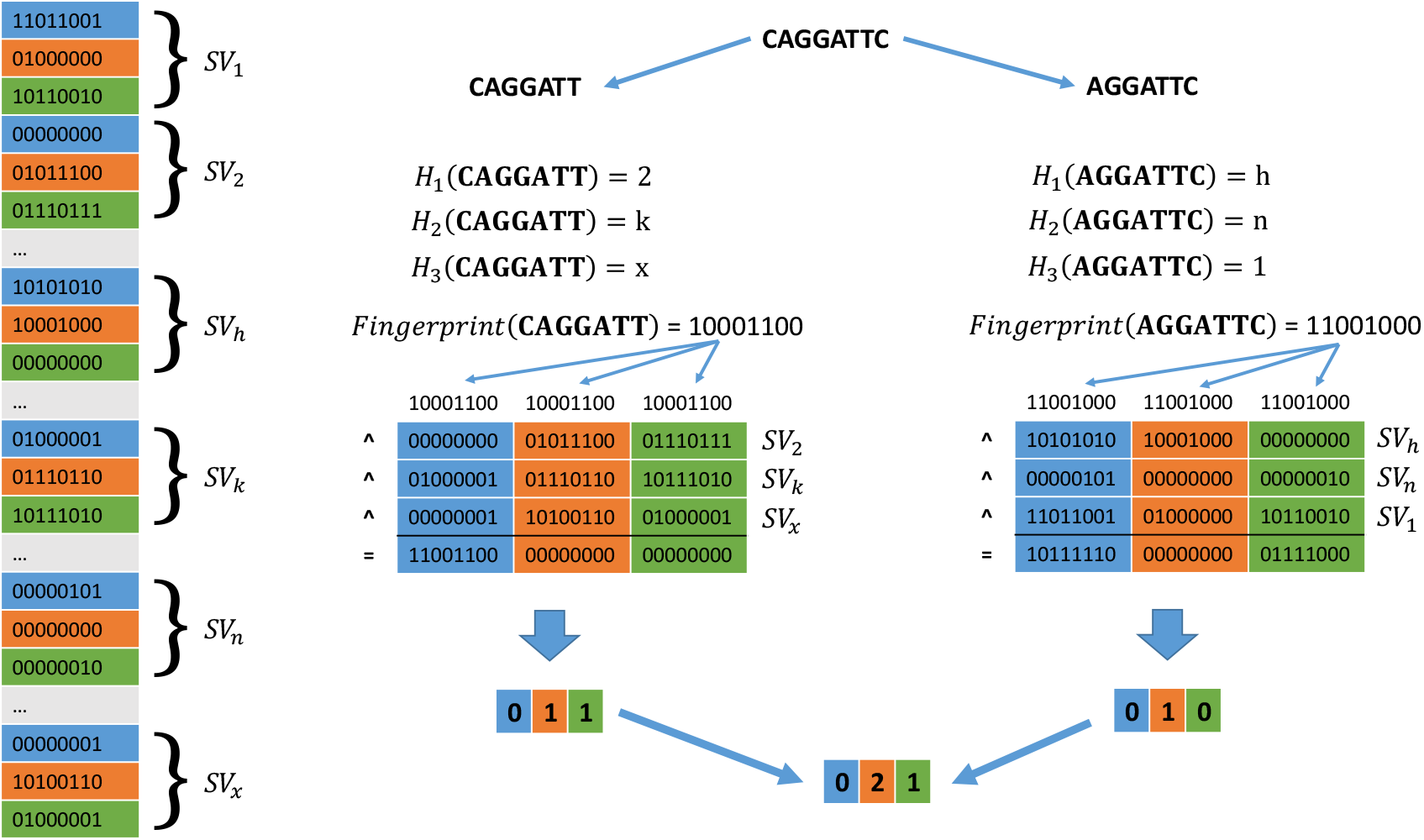
Querying an Interleaved XOR Filter consisting of three reference sequences. The read sequence CAGGATTC is divided into two overlapping 7-mers, CAGGATT and AGGATTC. For both k-mers, we separately calculate three hash values by applying the same hash functions used for building the IXF. For example, the hash values of k-mer CAGGATT point us to the subvectors *SV*_2_, *SV*_*k*_, and *SV*_*x*_. We further calculate the fingerprint for CAGGATT and concatenate the resulting 8-bit sequence three times because the IXF stores three reference sequences, which results in each subvector having three bins. Applying a bitwise XOR to the three subvectors and the fingerprint vector results in an 8-bit sequence for each reference. For k-mer CAGGATT, the second and the third bin equal to 00000000, which indicates that the k-mer is present in the second and third reference. This information is stored in a binning bitvector, e.g., 011 for k-mer CAGGATT. Finally, the binning bit-vectors of both k-mers are combined into a counting vector that stores the number of k-mer matches between the read and each reference sequence. Here, the vector 021 indicates that both k-mer are present in the second reference, and thus the read could match that reference.

**Fig. 5.**
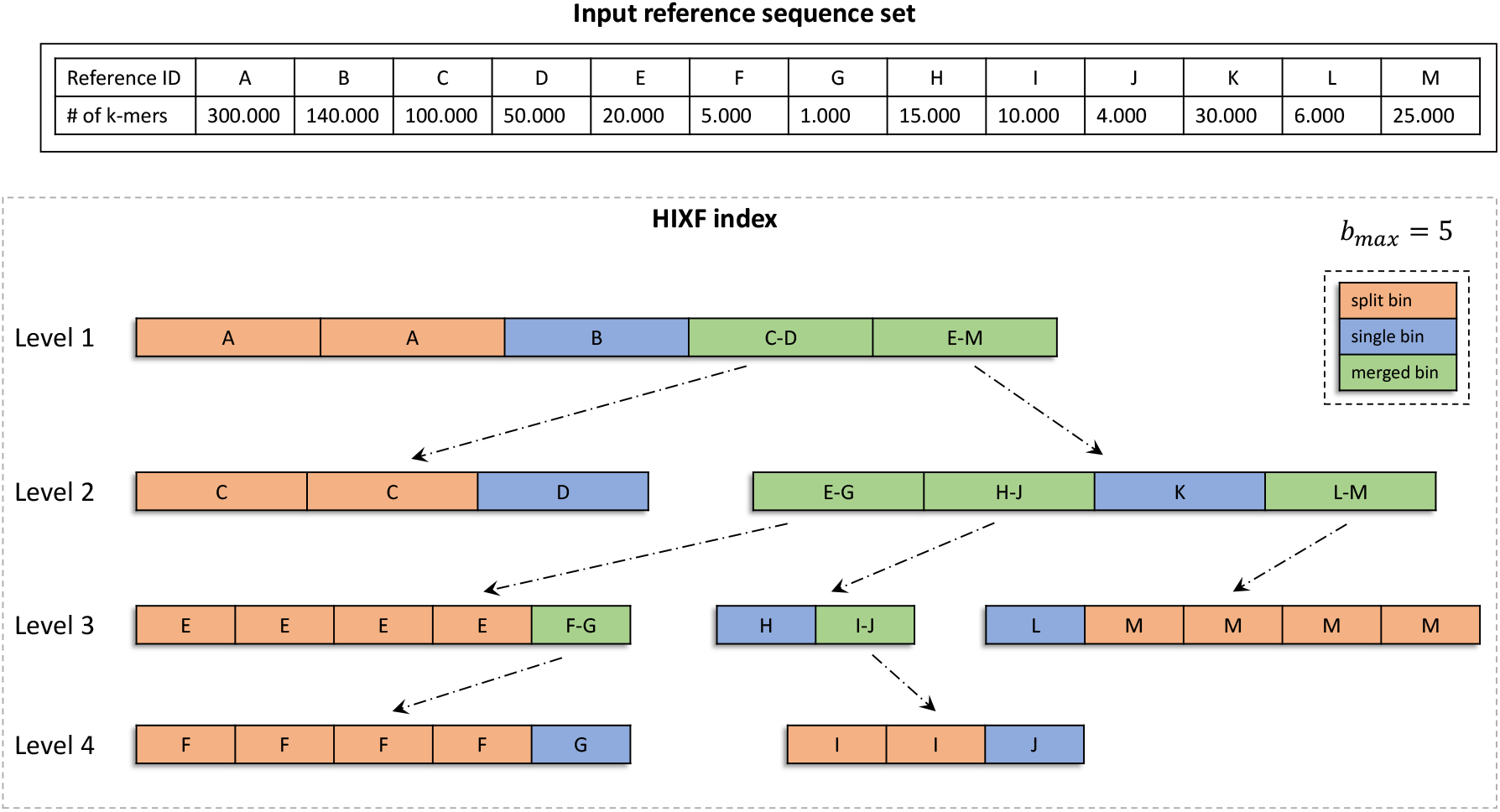
Building a HIXF index structure from 13 reference sequences. Exemplary building of a hierarchical interleaved XOR filter from thirteen differently sized reference genomes. Depending on the size of the input sequence’s k-mer sets, the HIXF layout is computed. Here, the maximum number of bins per IXF *b*_*max*_ = 5, which results in 4 levels consisting of 8 interleaved XOR filters. The first level is always a single IXF with exactly *B*_*max*_ bins. In the figure, Level 1 consists of two split bins for reference A, one single bin for reference B, one merged bin that stores all k-mers from references C and D, and one that contains the k-mer content of references E to M. On the second level, there is one additional IXF for each merged bin on the first level, resulting in two IXFs in the example above, where the first IXF consists of two split bins for reference C and one single bin for reference D. The second IXF on Level 2 consists of one merged bin containing all k-mers of references E, F and G, one additional merged bin with the k-mer content of references H, I and J, one single bin for reference K and another merged bin with the combined k-mer content of references L and M. Merged bins on each level of the HIXF result in an associated IXF on the next lower level forming a tree-like layout, where the leaves only have split and single bins.

**Fig. 6.**
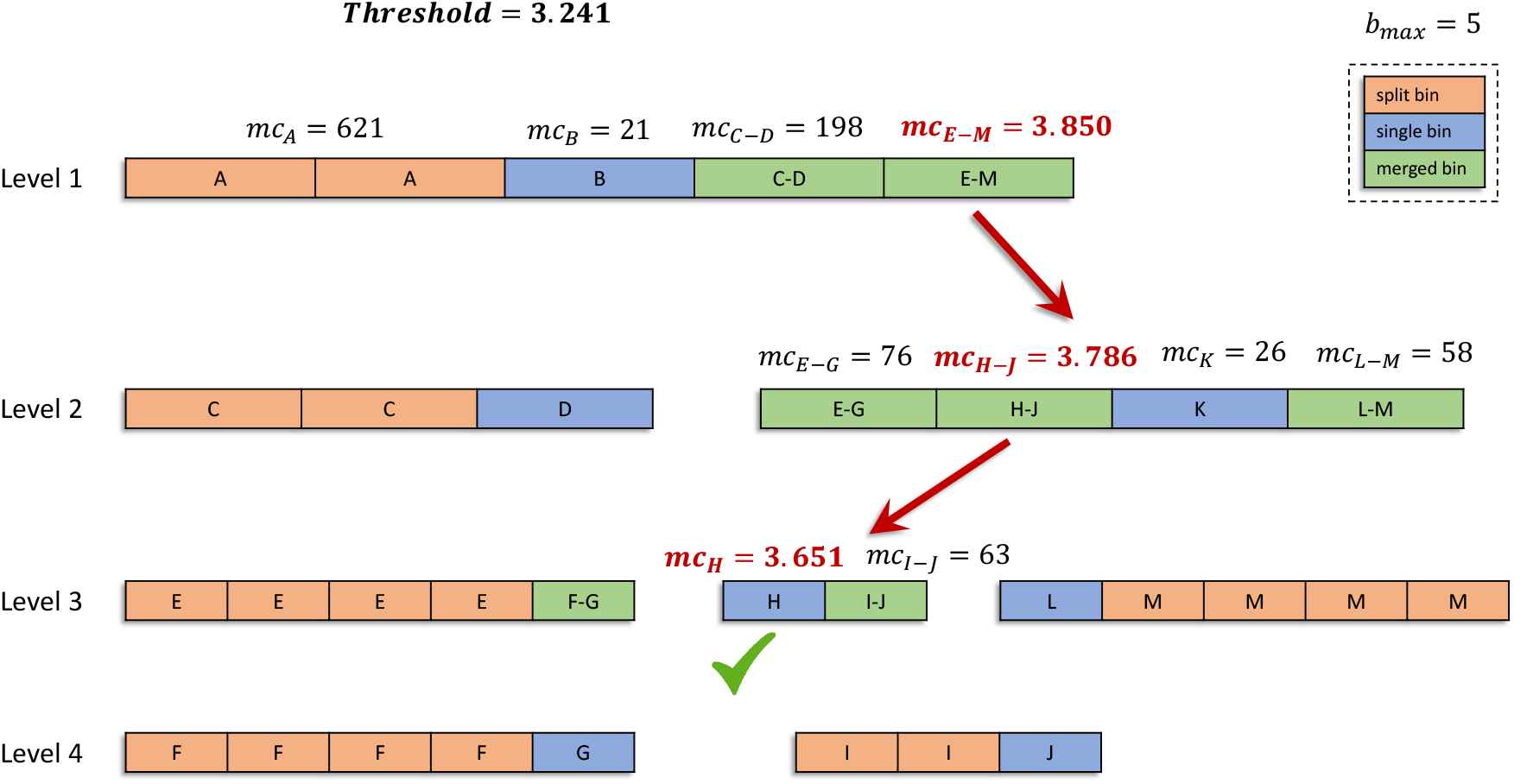
Querying a HIXF index structure of 13 reference sequences. When finding matching reference sequences for a given sequencing read, the k-mer content of the read is calculated first. Based on the expected sequencing error rate, a threshold for the minimum number of found k-mers is determined to consider a reference sequence as a hit. Then, the number of matching k-mers with each bin is computed for the first level IXF, considering summed match counts of split bins and match counts of single bins that exceed the threshold as a hit. If the match count for a merged bin exceeds the threshold, we query the associated IXF on the second level. All levels of the HIXF are queried recursively until all reference sequences with k-mer match counts exceeding the threshold have been found. In this example, only the match count for the merged E-M bin exceeds the threshold on the first HIXF level, and we continue querying the associated IXF on the second level. Here, only the merged H-J bin has a k-mer match count above the threshold, which requires querying the associated IXF on the third level, where we find the single bin of reference sequence H to exceed the match count threshold only. Thus, we report reference H as a hit with the queried sequencing read.

### Hierarchical Interleaved XOR Filter

The interleaved nature of the IXF has two important limitations. First, the largest XOR filter determines the overall size of the IXF because all single bins (XOR filters) of the IXF must have the same size. This means that the largest reference sequence dictates the size of the XOR filters storing the k-mer contents of the other reference sequences. Consequently, we would waste a substantial amount of space for smaller references if the reference sequences in the IXF have highly divergent sizes. Second, the query speed slows down with an increasing number of XOR filters stored in the IXF. This is, in practice, not a problem for a few hundred to a few thousand references, but becomes inefficient when storing many thousands of reference genomes.

To overcome this issue, we adapted the approach by Mehringer et al. to create a hierarchically structured interleaved XOR filter (HIXF). Here, the idea is to split the k-mer content of larger reference sequences into several smaller k-mer sets while merging the k-mer sets of very small reference sequences into one big set of k-mers. The resulting k-mer sets are stored in a high-level IXF, and for each merged k-mer set, a low-level IXF is stored, holding the k-mer sets of the smaller reference sequences in individual bins (XOR filters). While splitting the k-mer content of large reference sequences and merging the k-mer content of smaller references avoids wasting space, recursively adding an IXF for each merged k-mer set enables querying the individual k-mer sets of the small reference sequences. Depending on the number and size of the reference sequences and the maximum number of bins allowed for each IXF, we can have many levels of the hierarchical interleaved XOR filter (HIXF).

To compute the layout of the HIXF, we utilize the dynamic programming (DP) approach from Mehringer et al. that finds the optimal balance between space consumption and query speed. Here, we first calculate HyperLogLog sketches (38) to determine the Jaccard distance between each pair of reference sequences (39). Based on this information, the list of reference sequences is rearranged to specify which reference k-mer sets are more similar and, thus, ideal targets for merging on higher levels of the HIXF. Then, the dynamic programming algorithm uses a scoring function accounting for the space consumption of each IXF and the number of levels in the HIXF, which finds the optimal splitting and merging operations for the given set of references. Finally, a score is calculated for different values of the maximum number of XOR filters *b*_*max*_ in the IXFs, accounting for the expected query time and memory consumption. In general, *b*_*max*_ is a multiple of 64, and we calculate the score until the product of query time and space requirement increases. The minimum score determines the maximum number of XOR filters in each IXF, and the backtracing in the corresponding DP matrix returns the layout of the HIXF.

When querying the HIXF, we first calculate the k-mer content of the given query read and determine the minimum number of matching k-mers with each reference sequence to be considered a hit. We calculate this threshold based on the k-mer selection scheme described in the following subsection. Then, for each k-mer of the read, the membership in all filters of the top-level IXF is determined, and we further count the total number of k-mers that match each bin in the top-level IXF. If the counter for a bin exceeds the calculated threshold, the read is considered a match with the corresponding reference sequences. For the split bins, the k-mer counts have to be accumulated before thresholding, while we can directly answer the query for single bins that are neither merged nor split. For merged bins that exceed the threshold, we apply the same procedure recursively on the associated child IXF on the next lower level. This approach allows us to skip querying all lower-level IXFs whose upper-level merged bins do not exceed the given threshold. As a final result, we obtain a list of all reference sequences that exceed the minimum number of matching k-mers with the read under investigation.

### K-mer selection & thresholding

In the previous subsections, we introduced our new HIXF data structure describing the usage of membership queries for all k-mers of a given read against all k-mers of a given reference sequence set. We consider a read a hit with a reference sequence if the number of matching k-mers is greater than or equal to a given threshold *t*. For this k-mer model, we calculate the threshold as described in the Supplemental Material.

Since using all k-mers of the reference sequences can result in huge index sizes, k-mer selection approaches gained much attraction during the last decade, with minimizers being the most popular down-sampling approach for metagenomic classification of short reads (4, 16). However, Edgar recently showed that syncmers are more sensitive for selecting conserved k-mers in biological sequences and Dutta et al. could also improve long-read mapping by using open canonical syncmers instead of minimizers. Therefore, we implemented open canonical syncmers as a down-sampling strategy for large sets of k-mers to decrease the size of the HIXF index. In our implementation, open syncmers are sampled based on three parameters (*k, s, t*) where *k, s*, and *t* are positive integers and *s* ≤ *k*. The method then compares the *k*−*s* + 1 consecutive s-mers within a k-mer and selects the k-mer as a syncmer if the smallest s-mer occurs at position *t*∈ [0, *k − s* + 1] within the k-mer. The smallest s-mer is defined by the hash value computed for each s-mer. We use the canonical representation of syncmers, meaning that the lexicographically smallest syncmer out of its forward and reverse-complement sequence is always selected.

Analogous to the k-mer-based approach, we need to determine a threshold for a read’s minimum number of matching syncmers to consider it a hit with a reference sequence. In contrast to k-mers, there is no theoretical derivation for a (1 − *α*) confidence interval of the number of erroneous syncmers. Thus, we decided to derive the threshold empirically by simulating error-prone nanopore reads from a random sample of 1,000 bacterial reference genomes from the GTDB (19). We used the Rust implementation of the read simulator Badread (40) to simulate nanopore reads with five different read lengths between 1,000 and 5,000 bp and repeated the simulations for 20 different read accuracy rates (80%, 81%, 82%, …, 99%). Next, we build separate HIXF index files of the 1,000 genomes, one for each even k-mer value between 16 and 30. We only allow for even-numbered values for the k-mer size because we use canonical syncmers, with 16 being the practically smallest value to distinguish k-mers from different reference sequences and 30 the maximum value that allows finding k-mer matches between error-prone reads and a reference sequence. Finally, we separately queried the simulated reads of different read accuracies against the created HIXF index files and calculated the minimum fraction of found syncmers for each read and every combination of read accuracy and k-mer size. This minimum matching ratio for the different read accuracy and k-mer size combinations is used by Taxor to calculate the threshold for the minimum number of syncmer matches between a given read and the reference sequences stored in the HIXF index.

### Taxonomic profiling

The existence of homologous regions of genome sequences across multiple microbial species can lead to high false positive rates if taxonomic profiling methods exclusively rely on sequence similarity information. However, setting a low sequence similarity threshold is essential for detecting all species in a sample if the sequencing accuracy hardly reaches values of 98%. Therefore, we apply a three-step filtering approach of potential hits between reads and reference sequences before refining the results using an expectation-maximization (EM) algorithm that re-assigns reads to references based on the number of k-mer matches and taxonomic abundances of matched references.

Before the filtering, reads are assigned to matched reference genomes if the number of matching k-mers exceeds a certain threshold. Thus, a read can be assigned to many reference genomes in the index. We perform the first filter step on the single read level, determining the best matching reference genome based on the maximum number of k-mer matches (*max*_*kmatch*_) with the given read. All reference assignments to that read where the number of matching k-mers is smaller than 0.8× *max*_*kmatch*_ are considered spurious matches and removed from the results. The second filtering step creates a list of all reference genomes with at least one uniquely mapped read (a read assigned to exactly one reference genome). Consequently, we remove all read-to-reference assignments in the results where the reference genome has no uniquely mapped read. Finally, we apply the two-stage taxonomy assignment algorithm used in MegaPath (41) to reduce suspicious matched references. In short, the algorithm identifies reference genomes with less than 5% of their matches being uniquely matched reads. If such a reference genome *S* also shares a certain amount (e.g., 95%) of matches with another reference genome *T*, all matches of reads to *S* are re-assigned to *T*.

After filtering, we estimate the relative abundances of all matched references and reassign reads using a standard EM algorithm. This approach iteratively maximizes the likelihood that a given read *r* comes from a reference genome *g*, which also maximizes the likelihood of the relative taxonomic abundances in the whole read set. Let *G* be the set of genomes in the database and let *π*_*g*_ be the probability that a sequencing read in the sample emanates from database genome *g* ∈ *G*. We define the likelihood of the mapped read set *R* as

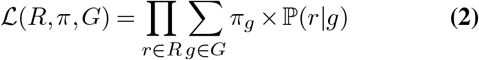

where ℙ (*r*|*g*) is the probability of read *r* coming from reference genome *g*. We use *mc*(*r, g*), the number of k-mer (or syncmer) matches between read *r* ∈ *R* and genome *g* ∈ *G*, and define ℙ (*r*|*g*) as

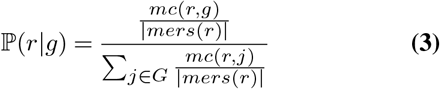

with |*mers*(*r*)| being defined as the number of k-mers (or syncmers) computed from *r*. After initialization of the axonomic compositions with 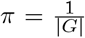 for all *g* ∈ *G*, we calculate in each iteration step an updated read assignment for each *r* ∈ *R* by

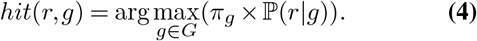

Based on the reassignment of reads, we update the taxonomic compositions *π* in each iteration step by accumulating the read lengths of all reads mapping to a certain genome and normalizing this value by the genome length.

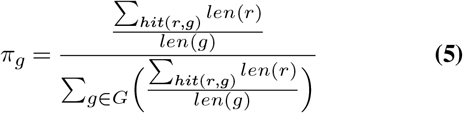

The nominator in Equation 5 can be interpreted as the depth of coverage on genome *g* in the sample under investigation. We divide the coverage of *g* by the sum of all genome coverages to get the relative taxonomic abundance of *g* in the sample. The single steps of the EM algorithm are repeated until convergence of the likelihood ℒ (*R, π, G*) or after a predefined number of iteration steps (default 10).

## Results

We compare Taxor to five state-of-the-art taxonomic profiling tools, Centrifuge (3), MetaMaps (5), Kraken2 (4), KMCP (17) and Ganon (16). Centrifuge is the tool underlying Oxford Nanopore’s “What’s in my pot” (WIMP) application for real-time species identification (42). While Centrifuge was initially developed to analyze short-read metagenomic samples, MetaMaps specifically addresses the task of strain-level metagenomic assignment of long reads. We decided to use both tools and Kraken2 with default parameter configurations because this results in the best recall and precision in our experiments on simulated data.

Finally, we evaluated KMCP and Ganon, both utilizing different Bloom Filter approaches to store sets of selected k-mers from reference genomes for short-read classification. For Ganon’s classify module, we have to set parameters “–rel-cutoff 0.12” and “-rel-filter 0.9” to account for the higher error rates in nanopore reads. For the same reason, we also set parameters “–min-query-cov 0.12”, “–min-hic-ureads-qcov 0.2” and “–min-chunks-fraction 0.2” when running KMCP’s search and profile subcommands. For the evaluation of Taxor, we only set the parameter “–error-rate 0.15”, trying to assign all nanopore reads with error rates smaller than or equal to 15%. The specific commands to run the six tools are provided in the Supplementary Data.

### Reference databases

Most taxonomic profilers offer prebuilt reference databases, but also allow building custom reference databases. Since the choice of the reference database directly affects the outcome of taxonomic profiling, a fair comparison between tools also requires using the same database. This ensures that observed differences in the single-read assignments are attributed solely to the profiling methods. Thus, we downloaded all complete genome sequences and chromosomes of archaea, bacteria, viruses, and fungi from the NCBI RefSeq database (Release 217) (18) using genome_updater 0.5.2 (https://github.com/pirovc/genome_updater). We used only one reference genome per species, resulting in 21,003 genomes used to build custom databases for each tool. This custom database consists of 11,579 viral genomes, 8,938 bacterial genomes, 403 archaea genomes, and 83 fungi genomes.

We build customized index data structures based on the described reference database for all tools included in our evaluation. For Centrifuge, we created the reference index using the default parameters, providing only the taxonomic information from the NCBI taxonomy and reference sequences in fasta format. For MetaMaps, we first created a custom database from our downloaded taxonomy using the provided scripts and following the instructions on https://github.com/DiltheyLab/MetaMaps. Then we created the MetaMaps index from the custom database using default parameters. Since the other tools all use pseudo-alignment utilizing k-mer-based approaches, we build indexes using a k-mer size of 22, which is a good compromise between high specificity on species-level identification and high sensitivity for error-prone nanopore read classification. For all four approaches, we decided to use k-mer selection schemes that downsample the used k-mer sets to roughly 10% of all reference k-mers to reduce memory usage and index size significantly. Specifically, we used ungapped k-mers of size 22 and a window size of 32 for minimizer-based indexes in Kraken2 and Ganon. For KMCP and Taxor, we used a k-mer size of 22 and a syncmer size of 12. We further set the false positive rate for the Bloom Filter-based approaches, namely Ganon and KMCP, to 0.3% to reflect the same inherent false positive rate of Taxor’s XOR filter approach. The specific commands and instructions to build the reference indexes of all tools are listed in the Supplementary Data.

### Evaluation datasets

We carried out four experiments to evaluate Taxor, covering multiple metagenome composition scenarios from simulated and real data as well as sample contamination with eukaryotic DNA. The simulated dataset consists of 100 randomly chosen reference genomes included in the reference database and comprises 1,124,128 reads. For read simulation, we used pbsim2 (43) with parameters “–accuracy-mean 0.95”, “–length-min 1000” and “–hmm_model R103.model”. We chose a mean accuracy of 95% because the latest basecallers for ONT data have shown to reach such accuracies (44). We further decided to simulate reads with a minimum read length of 1,000 because shorter nanopore reads are commonly misclassified, and thus, MetaMaps and Taxor do not classify those reads.

For evaluation on real data, we obtained two ONT datasets for the ZymoBIOMICS D6300 microbial community standard (45) and one PacBio HiFi dataset for the ZymoBIOMICS Gut Microbiome Standard D6331. The Zymo D6300 standard consists of ten evenly abundant species, including 8 bacteria at 12% sequence abundance and two yeasts at 2% sequence abundance. The first ONT dataset comes from a continually updated online resource (https://lomanlab.github.io/mockcommunity/r10.html). We downloaded the R10.3 chemistry data release (February 2020), which was produced from two flowcells on an ONT GridION, resulting in 1.16 million reads (4.64 Gb data). The second ONT dataset was obtained from the European Nucleotide Archive (PRJEB43406: ERR5396170, released March 2021) and represents the ‘Q20 chemistry’ release for the Zymo D6300 standard (described at github.com/Kirk3gaard/2020-05-20_ZymoMock_Q20EA). It was generated using a PromethION, resulting in 5.4 million reads (17.95 Gb data).

The PacBio HiFi dataset for the ZymoBIOMICS Gut Microbiome Standard D6331 (PRJNA680590: SRX9569057, released November 2020) contains 17 species (including 14 bacteria, one archaeon, and two yeasts) in staggered abundances. Five species occur at 14% sequence abundance, four at 6%, four at 1.5%, and one per 0.1%, 0.01%, 0.001%, and 0.0001% abundance level. There are five strains of E. coli in this community (each at 2.8% sequence abundance), which we treat here as one species at 14% sequence abundance. The PacBio Zymo D6331 dataset was generated using the Sequel II System and contains 1.9 million HiFi reads with a median length of 8.1 kb, for a total of 17.99 Gb of data.

To generate a read-level truth set, we use minimap2 (46) to map the reads against the reference genomes provided by ZymoBiomics. All reads that cannot be mapped with minimap2 are excluded, and the primary alignment for each read determines the assumed true placement. This results in two ONT Zymo D6330 evaluation datasets referred as “ZymoR10.3” and “ZymoQ20”, and one PacBio HiFi Zymo D6331 evaluation dataset referred as “HiFi_D6331”.

As a negative control, we use pbsim2 with the same parameters described above to simulate long-read sequencing data from two eukaryotic genomes not present in the reference database. Specifically, we simulate 685,303 reads from the *Aedes aegypti* (yellow fever mosquito) genome (GCF_002204515.2) and 142,677 reads from the *Toxoplasma gondii* ME49 genome (GCF_000006565.2). The two read sets are analyzed independently with Taxor and the other tools.

### Evaluation metrics

We evaluated the performance of all six tools using several criteria. We assessed read utilization and classification metrics at the species, genus, and family levels and relative abundance estimates at the species level. First, we evaluated read utilization for each profiling method by calculating the total percent of reads assigned to specific taxonomic levels. We performed this for the following ranks: class, order, family, genus, and species. Here, we expected methods like Kraken2 and Ganon that use an assignment to the lowest common ancestor to display read assignments across multiple taxonomic levels, while methods like Taxor only report the species level.

We calculated several metrics to evaluate read classification performance based on the number of true positives, false positives, and false negatives. In this context, we define a true positive as a correct taxon assignment of a read. We define a false positive as an incorrect taxon assignment on the read level. We further define a false negative as the failure to detect a taxon for a specific read. The formulas for precision, recall, and F-scores are as follows:

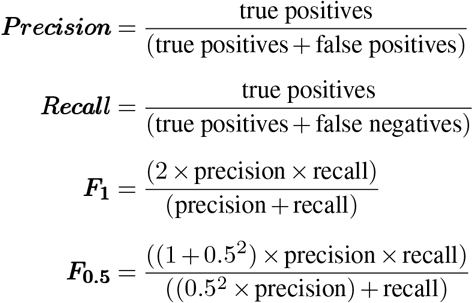

The values for the above metrics each range from 0 to 1. For precision, a score of 1 indicates that all reads with a taxon assignment have been assigned to the correct taxon, whereas lower scores indicate a higher number of wrong taxon assignments. For recall, a score of 1 indicates that all reads were assigned to the correct taxon, whereas a lower score indicates that no taxon could be assigned for some reads. The F-scores provide a useful way to summarize the information from precision and recall. The *F*_1_ score is the harmonic mean of precision and recall (both measures are weighted equally), whereas the *F*_0.5_ score gives more weight to precision (placing more importance on minimizing false positives). A value of 1 for either F-score indicates perfect precision and recall.

For accurate measurement of the read classification metrics for the real mock datasets, we had to control for species synonymies in the taxonomy of the used reference database. To avoid a negative impact on the metrics, we used the sum of cumulative counts for the species and all synonyms as the read count for the taxon. In particular, this included five species in Zymo D6300 (*Limosilactobacillus fermentum* = *Lactobacillus fermentum*; *Bacillus subtilis* = *Bacillus spizizenii*; *Escherichia coli* = *Escherichia sp. TC-EC600-tetX4*; *Listeria monocytogenes* = *Listeria sp. LM90SB2*; *Staphylococcus aureus* = *Staphylococcus sp. T93*) and two species in Zymo D6331 (*Limosilactobacillus fermentum* = *Lactobacillus fermentum*; *Escherichia coli* = *Escherichia sp. TMG17TGC*), where we treated the 5 strains of *E. coli* contained in this community as one species.

We calculated detection metrics for each dataset. To understand the performance of each method across all datasets, we took an average of precision, recall, *F*_1_, and *F*_0.5_ at species level for the simulated and real datasets.

Finally, we attempted to obtain relative abundances for each method, acknowledging differences in reporting abundances as described by Sun et al. In particular, there are clear differences in intended outputs among methods. For example, profiling methods (Ganon & KMCP) provide taxonomic abundances, whereas classifiers (Centrifuge, Kraken2 & MetaMaps) provide sequence abundances. Since Taxor reports both abundance measurements, we did not transform the reported values of the tools but compared Taxor’s output directly to the reported abundance types of the respective tools. We calculated an L1 distance between observed and theoretical abundances for each method as described by Portik et al. The theoretical abundances were obtained from the manufacturer’s specifications based on genomic DNA (sequence abundance) and genome copy (taxonomic abundance). We calculated the L1 distance by summing the absolute error between the theoretical and empirical estimate per species across the three communities. In this calculation, we included the false positives lumped in the “Other” category and compared them against a theoretical abundance of zero for this category.

### Read utilization performance

Across the four metagenomic evaluation datasets, Taxor shows a high total read assignment between 89% (ZymoR103) and 97% (Hifi_D6331). Since we did not implement a lowest common ancestor algorithm in our new tool, all reads have been directly assigned to the species level. Compared to the other five tools in our benchmarking (see Fig. 7), Taxor generally classifies fewer reads than Centrifuge, Kraken2, and MetaMaps but more than KMCP and Ganon. However, for the real datasets, Taxor assigns more reads than Kraken2 and Centrifuge at the species level. Both tools assign a considerable amount of reads to the genus level, particularly for the ONT datasets. KMCP shows by far the lowest total read assignments of all tools on the ONT datasets. For the PacBio HiFi_D6331 dataset, all tools show a comparable high percentage of read assignments on the species level (between 92% for Kraken2 and 97% for Taxor and Ganon), except for KMCP, which assigns 67% to species level, 12% to genus level and 16% to family level. In general, we observe that the read utilization of some tools is highly dependent on the dataset and especially the underlying sequencing technology, with ONT data having significantly higher error rates than PacBio data.

**Fig. 7.**
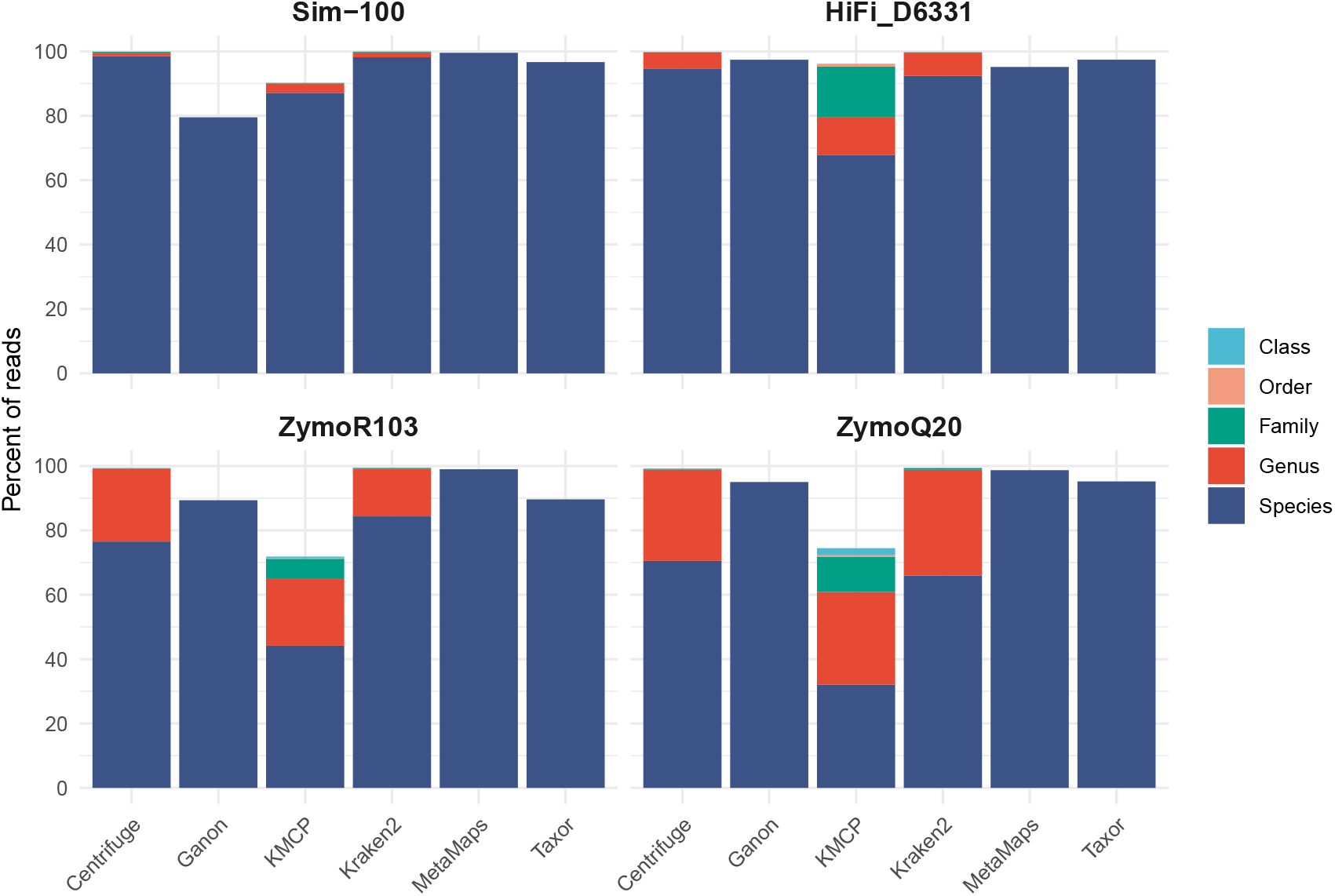
Read utilization of simulated and real metagenomic datasets. The stacked barplots show the total percent of reads that were assigned to different taxonomic ranks, highlighted in different. Taxor generally classifies fewer reads than Centrifuge, Kraken2, and MetaMaps but more than KMCP and Ganon. For the real datasets, Taxor assigns more reads than Kraken2 and Centrifuge at the species level.

### Classification performance on simulated data

We first evaluate the read classification performance of Taxor in a simulation experiment, which represents a medium-complexity metagenomic analysis scenario with 100 randomly chosen species from the used reference database. We report the resulting performance metrics on the species level for all six evaluated tools in Table 1. On this dataset, Taxor read assignments achieve a recall of 0.96, a precision of 0.99, and F-scores of 0.98 and 0.99. Taxor outperforms KMCP and Ganon in terms of recall by 9-17% while having slightly lower recall than Centrifuge, MetaMaps, and Kraken2 (2-3%). All tools show a high read classification precision in this experiment ranging from 0.98 to 0.99, which means the tools report very few false read assignments. Four of the six tools can also correctly classify more than 95% of the simulated reads. Only Ganon and KMCP show a lower recall, failing to classify 13-21% In a second simulation experiment, we assess the effect of contamination with eukaryotic host DNA from larger genomes on read classification. Therefore, we simulated reads from two eukaryotic genomes (*Aedes aegypti* and *Toxoplasma gondii*), neither of which is present in the reference database. Taxor has a low false-positive rate for both read sets and correctly leaves the large majority of reads unclassified (≥99.99%) on all taxonomic levels. We also note low false positive rates for KMCP (0% for both datasets) and Ganon (0.24% for *Aedes aegypti* and 0.05% for *Toxoplasma gondii*). The three tools slightly outperform MetaMaps, having misclassification rates between 1.71% (*Aedes*) and 2.87% (*Toxoplasma*). In contrast, Kraken2 and Centrifuge show high false-positive rates, with Kraken2 reporting only 5.67% (*Aedes*) and 8.84% (*Toxoplasma*) of reads as unclassified, and Centrifuge reporting 22.90% (*Aedes*) and 26.74% (*Toxoplasma*) of reads as unclassified. Detailed results for these experiments are provided in Supplementary Table S3.

**Table 1.**
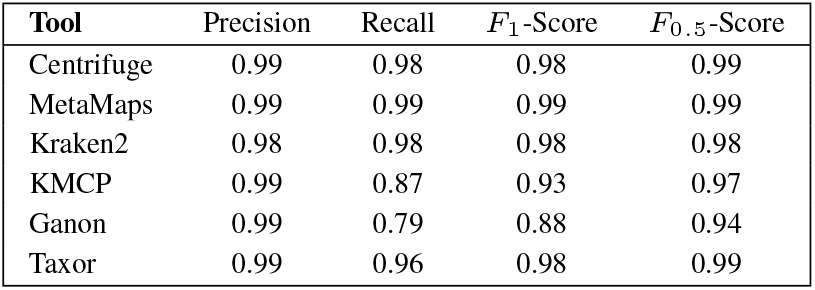
Species-level read classification performance on simulated data. Precision, recall, and F-scores for the species-level analysis of simulated Data. All compared tools show high precision and recall between 0.96 and 0.99, except for Ganon and KMCP, which have a lower recall of 0.79 and 0.87, respectively.

| Tool | Precision | Recall | $F_1$-Score | $F_{0.5}$-Score |
| --- | --- | --- | --- | --- |
| Centrifuge | 0.99 | 0.98 | 0.98 | 0.99 |
| MetaMaps | 0.99 | 0.99 | 0.99 | 0.99 |
| Kraken2 | 0.98 | 0.98 | 0.98 | 0.98 |
| KMCP | 0.99 | 0.87 | 0.93 | 0.97 |
| Ganon | 0.99 | 0.79 | 0.88 | 0.94 |
| Taxor | 0.99 | 0.96 | 0.98 | 0.99 |

### Classification performance on real data

Since the simulated dataset represents only a medium-complex metagenomic sample, we further evaluate Taxor on real data using three sets of metagenomic sequencing data, two ONT datasets for the ZymoBIOMICS D6300 community standard, and one PacBio HiFi dataset for the ZymoBIOMICS Gut Microbiome Standard D6331. The read classification performance evaluation results are shown in Fig. 8 and Supplementary Table S2. Consistent with observations on simulated data, Taxor shows a high precision on the species level of 0.97 and 0.94 on the ZymoR10.3 and HiFi_D6331 data sets. For both datasets, The precision increases to 0.98 on the genus level and 0.99 on the family level. For the second ONT dataset (ZymoQ20), Taxor’s species-level precision is slightly lower at 0.91 and increases to 0.92 (genus) and 0.93 (family) on higher taxonomic levels. Taxor further shows a high recall between 0.95 and 0.97 on all taxonomic levels for the ZymoQ20 and HiFi_D6331 datasets. Only for the ZymoR10.3 data set, Taxor’s recall has a value of 0.89 on all taxonomic levels. Our new tool has a *F*_1_-Score between 0.93 and 0.96 and a *F*_0.5_-Score between 0.92 and 0.95 on the species level across all three datasets.

**Fig. 8.**
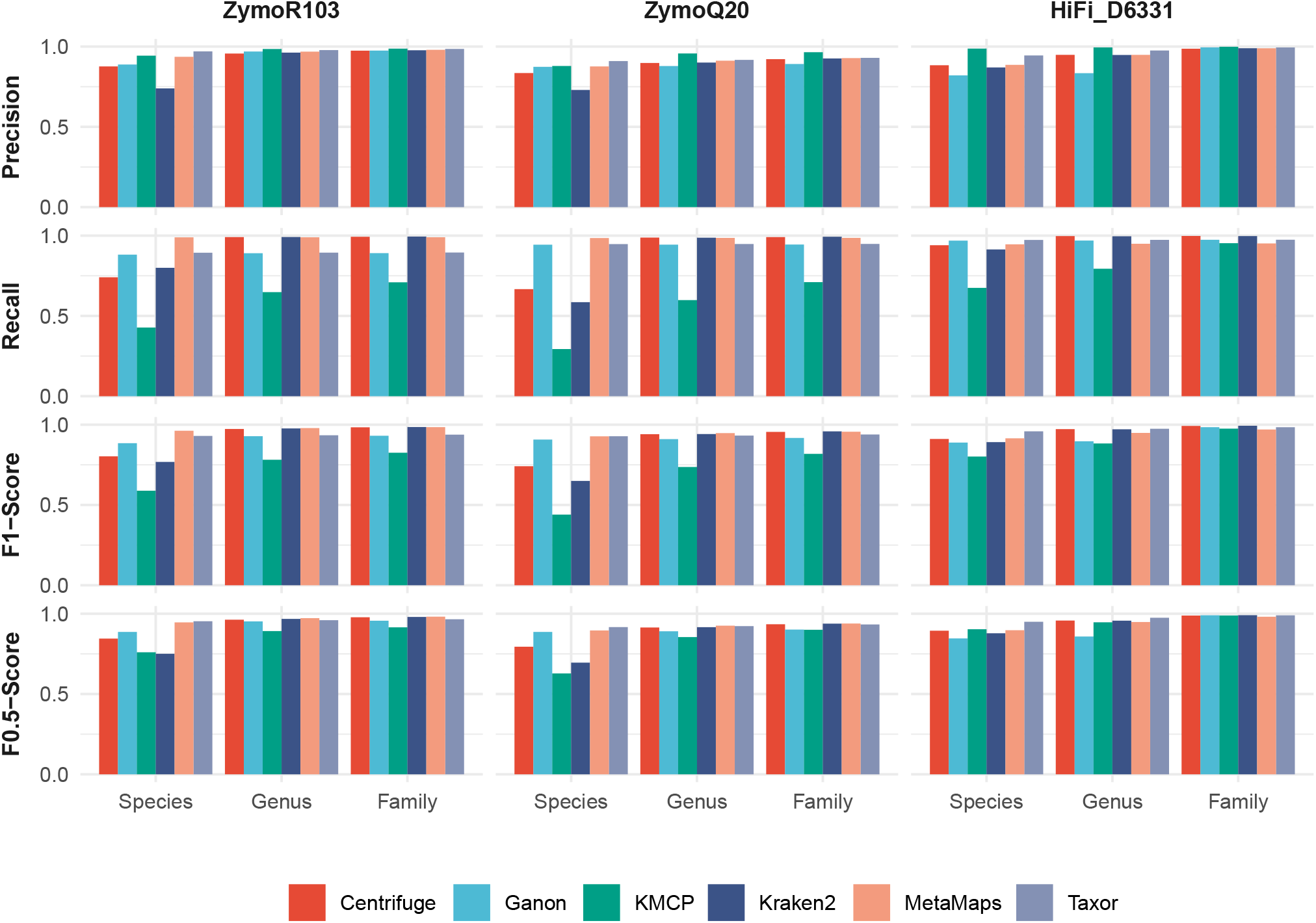
Read classification on species, genus, and family level for all three real datasets. Precision, recall, *F*_1_ -Score, and *F*_0.5_ -Score are shown for all six compared tools on three taxonomic levels. Taxor and KMCP show the highest precisions across all three datasets and all levels. Taxor’s recall is comparable to MetaMaps and Ganon on the species level, whereas KMCP consistently has the lowest recall of all tools. Centrifuge and Kraken2 have low species-level recall for the two ONT datasets while performing well on the HiFi dataset. Taxor and MetaMaps, in general, have the highest F-scores, with Taxor outperforming MetaMaps on the *F*_0.5_ -Score.

Consistent patterns emerge when comparing Taxor to the other metagenomics read classification tools. Across all taxonomic levels and all three datasets, Taxor and KMCP show the highest precision. On the species level, Taxor shows the highest precision on both ONT data sets (0.97 on ZymoR10.3 and 0.91 ZymoQ20), outperforming MetaMaps and KMCP by 1-3% and Ganon by 3-8%. On the HiFi_D6331 dataset, KMCP outperforms Taxor by 4% precision on the species level. However, KMCP shows the worst recall on all datasets across all taxonomic levels, while Taxor, on average, has the second-best recall on the species level (see Fig. 9). We further recognize that Centrifuge and Kraken2 have a low recall between 0.66 and 0.8 on the species level for both ONT datasets, which increases to 0.99 on the genus level. Since Taxor does not use a lowest-common-ancestor algorithm, its recall stays relatively constant across all taxonomic levels, which explains why Centrifuge and Kraken2 outperform Taxor in terms of recall on higher taxonomic levels. The long-read classification method MetaMaps has the highest average recall on the species level across the three datasets.

**Fig. 9.**
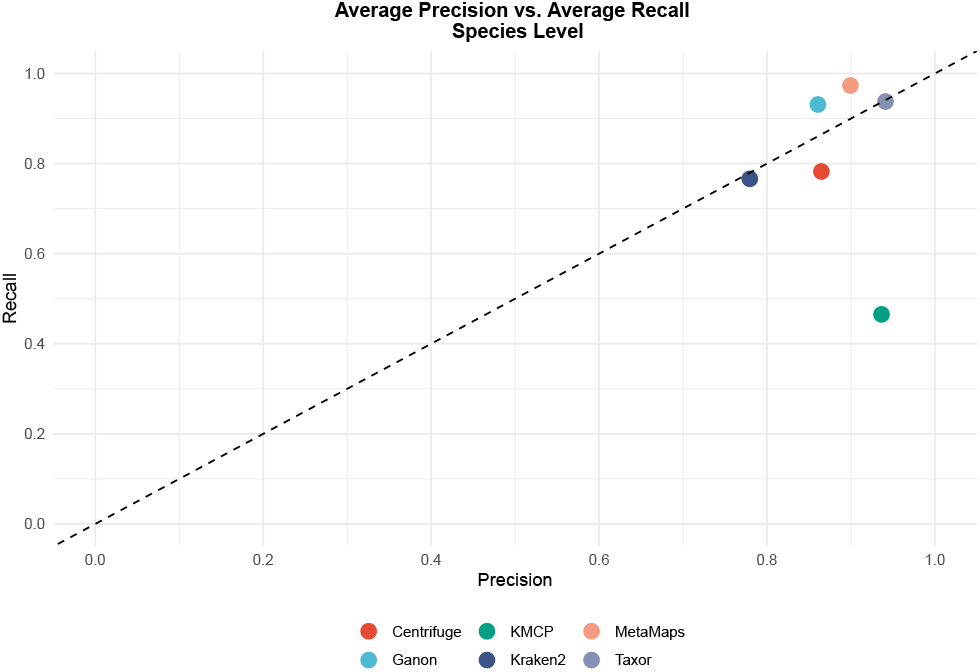
Average precision and recall on species level across the three real datasets. Showing average precision and recall of the six compared tools on the species level for the two ONT and the HiFi mock community datasets. MetaMaps, Taxor, and Ganon have the highest average recall, while Taxor and KMCP outperform the other tools in terms of precision. KMCP is the only tool with an average recall below 0.5 across the three datasets.

When looking at F-scores, we see that Taxor consistently outperforms the other tools on the species level, except for the *F*_1_-Score on the “ZymoR10.3” dataset, where MetaMaps performs best because of its high recall. On average, Taxor and MetaMaps have the same species-level *F*_1_-Score across all three datasets, but Taxor outperforms MetaMaps concerning the average *F*_0.5_-Score when precision is prioritized over recall (Fig. 10). On the genus and family level, all six tools are comparable except for KMCP, which suffers from low recall.

**Fig. 10.**
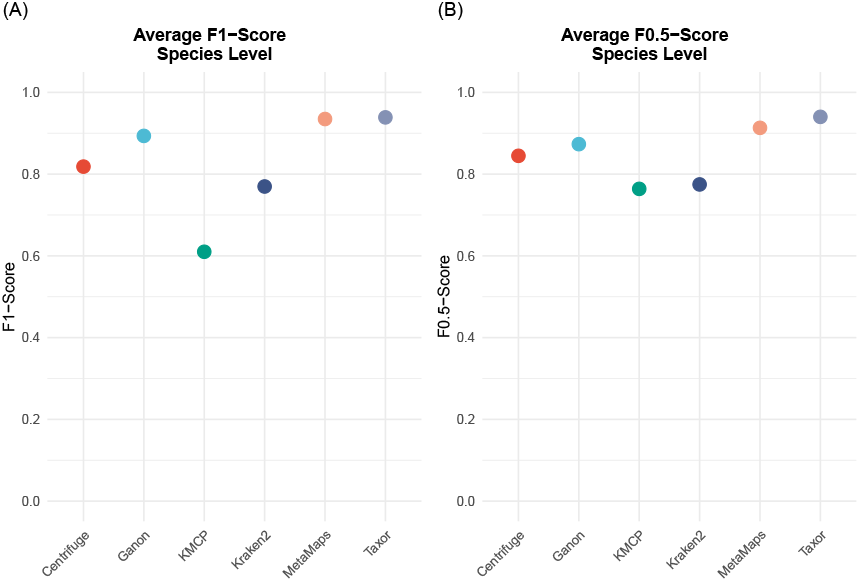
Average F-Scores on species level across the three real datasets. (A) Average species-level *F*_1_ -Score of the six tools across the three real mock communities. Taxor and MetaMaps show the highest scores, while KMCP has the lowest average *F*_1_ -Score. (B) Average species-level *F*_0.5_ -Score of the six tools across the three real mock communities. Our new tool Taxor outperforms the other methods when precision is prioritized over recall.

In summary, the two long-read methods, Taxor and MetaMaps, perform best on the species level across the three real datasets, making them the best choice for long-read metagenomics classification.

### Relative abundance estimation on real data

All six evaluated tools report relative species abundance estimations after read classification and metagenomic profiling. In Fig. 11, we compare the theoretical relative abundances of species in the mock communities against the relative species abundances reported by the different tools. Taxor is the only tool that reports both types of abundance, sequence abundance, and taxonomic abundance. When comparing the taxonomic abundance estimates of Taxor to those of Ganon and KMCP, we see that none of the three tools can accurately predict taxonomic abundances for all species in the three mock communities. However, Taxor has the lowest L1 distance of the three tools across all real datasets, demonstrating better relative species abundance estimates than Ganon and KMCP.

**Fig. 11.**
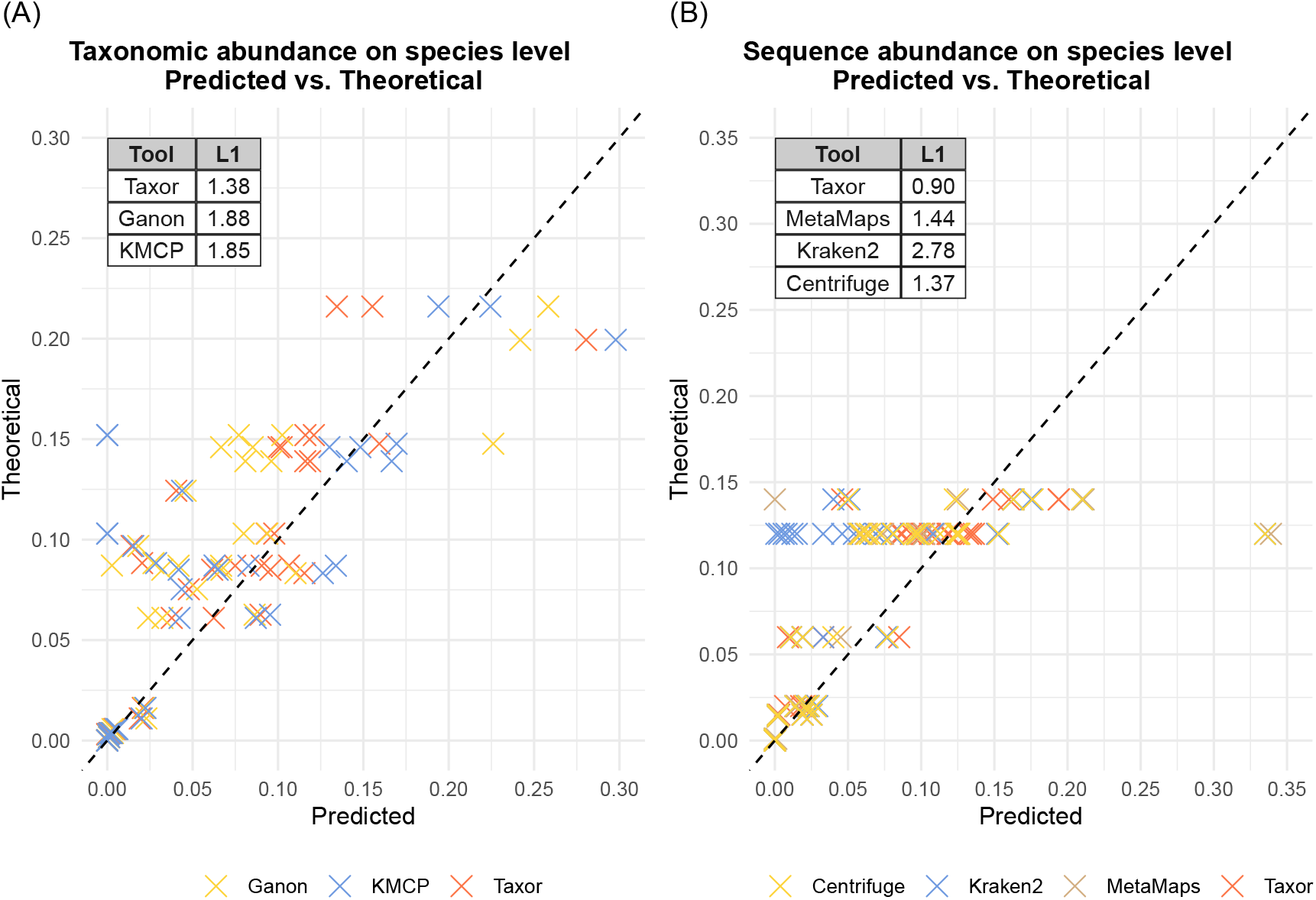
Comparison of theoretical abundances and predicted abundances. For each species in the three real mock communities, predicted abundances by the compared tools are plotted against the theoretical abundances. Crosses on the dashed line represent a perfect match between predicted and theoretical abundance. (A) Comparison of tools reporting taxonomic abundance. (B) Comparison of tools reporting sequence abundance. Taxor reports both abundance types and outperforms its competitors, having the smallest L1 distance between predicted and theoretical abundance.

Since MetaMaps, Kraken2 and Centrifuge report relative sequence abundances of species instead of relative taxonomic abundances, we compare their profiling results to the theoretical sequence abundances of the mock communities. Consistent with the taxonomic abundance evaluation results, none of the investigated tools can accurately predict sequence abundance for the species comprising the three mock communities. As for the taxonomic abundance, Taxor outperforms the other tools by having the smallest L1 distance between theoretical and predicted species abundances across all datasets.

### Computational requirements comparison

With the ever-increasing number of genomes in public databases comes the need for faster and more space-efficient methods to facilitate metagenomic read classification and profiling. Thus, we assessed the computational requirements of Taxor for indexing the reference database used in this study. Then we also measured the required peak memory usage and runtime for querying the “ZymoR10.3” dataset against the built database index. We further compare Taxor’s computational requirements against the other five tools used in this study. For all tools, we used the same reference database and built database indexes using the commands provided in the Supplementary Material. Specific commands for querying the built indexes are also provided in the Supplementary Material. We performed computational benchmarking on an AMD EPYC 7742 high-performance computing cluster (HPC) node using 30 threads. All times and peak memory usage were measured using the Linux “time” command with the parameter “-v”.

Results of the computational benchmarking are provided in Table 2. While Ganon is the fastest tool when building the index, this comes at the cost of high memory usage and a large index size on disk (≥120 GigaBytes). KMCP has the lowest peak memory usage (5.84 GigaBytes) and is almost as fast as Ganon, building the index in less than 10 minutes. Taxor and Kraken2 both need approx. 80 minutes to build an index, but need only 20% (Kraken2) and 10% (Taxor) of Ganon’s memory requirements. Although Taxor needs 2.5 times more memory than KMCP, the final HIXF index is 40% smaller than KMCP’s index and 65% smaller than the Kraken2 index. Centrifuge’s index is comparable to KMCP’s index size, but Centrifuge needs 344 minutes and 385 GigaBytes RAM to build its index. MetaMaps is the second resource-hungry tool in our benchmarking, needing three times as much time and 25 times as much memory as Taxor to build its index. MetaMaps further has the largest index size of all tools needing more than twice the disk space as Ganon and 25 times as much as Taxor.

**Table 2.**
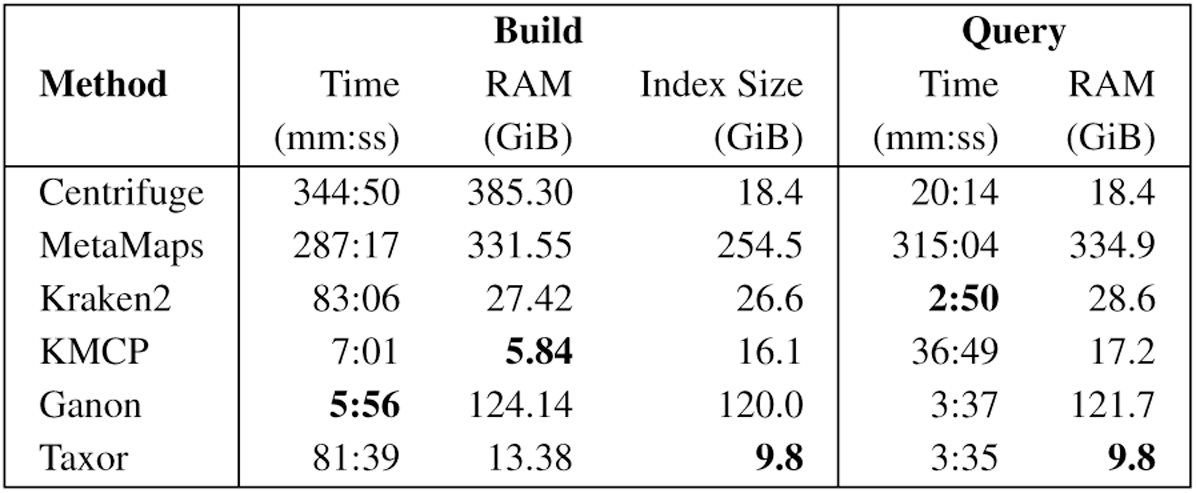
Results of Computational requirement benchmark. Reference indexes of a database consisting of 21,003 bacterial, viral, archaeal, and fungi genomes were built for all tools. We measured the elapsed time, peak memory usage, and index size for constructing the index. For the query benchmark, we measured the elapsed time and peak memory usage for classifying 426,213 ONT reads. Build and query times were measured using 30 threads on an HPC node.

| Method | Build |  |  | Query |  |
| --- | --- | --- | --- | --- | --- |
|  | Time (mm:ss) | RAM (GiB) | Index Size (GiB) | Time (mm:ss) | RAM (GiB) |
| Centrifuge | 344:50 | 385.30 | 18.4 | 20:14 | 18.4 |
| MetaMaps | 287:17 | 331.55 | 254.5 | 315:04 | 334.9 |
| Kraken2 | 83:06 | 27.42 | 26.6 | **2:50** | 28.6 |
| KMCP | 7:01 | **5.84** | 16.1 | 36:49 | 17.2 |
| Ganon | **5:56** | 124.14 | 120.0 | 3:37 | 121.7 |
| Taxor | 81:39 | 13.38 | **9.8** | 3:35 | **9.8** |

When querying the 426,213 ONT reads of the “ZymoR10.3” dataset against the respective database indexes, Kraken2 performs fastest, needing less than 3 minutes to report read classification results. Taxor and Ganon are almost as fast as Kraken2, with a query time of approx. 3 minutes and 30 seconds. These three tools outperform the others, with a six (Centrifuge) to 12 (KMCP) times faster query time. MetaMaps is the most resource-hungry tool, needing 335 GigaBytes memory and more than 5 hours to query all reads against its index. Finally, we note that Taxor outperforms all evaluated tools in terms of peak memory requirements when querying the index. Because of its small index size, Taxor can reduce the peak memory usage by approx. 50% compared to Centrifuge and KMCP, which both have much higher query times. Compared to Kraken2 and Ganon, which have similar query times, Taxor can reduce the memory footprint by a factor of three (Kraken2) to ten (Ganon). These results highlight the combination of fast and space-efficient read classification of Taxor, whereas other tools only provide fast querying or lower memory requirements.

## Discussion

The increasing size of public reference genome databases such as NCBI RefSeq makes metagenomic profiling and read classification a computationally challenging task. In particular, reducing memory usage has become an objective of many studies during the last few years. However, state-of-the-art shortand long-read metagenomic classifiers still consume large amounts of memory, which can only be reduced by accepting a higher risk of false classifications. For Bloom Filter approaches like Ganon and KMCP, one can, for example, reduce the index size by accepting a higher false positive rate for the approximate membership queries of k-mers. However, higher false positive rates can lead to false classifications of viral reads, as proposed by Shen et al. This issue motivated us to develop a new data structure that combines low memory requirements, a low false positive rate, and fast membership queries to facilitate precise metagenomic classification of long reads on large reference sequence databases. Based on the work of Graf and Lemire and Mehringer et al., we created a hierarchical interleaved XOR filter (HIXF) data structure, which is implemented in the metagenomic classification tool called Taxor. Instead of relying solely on k-mers, Taxor can utilize syncmers as a k-mer selection scheme. Finally, our new tool comes with a profiling step, applying several filter strategies and an EM algorithm that refines the results of the initial read classification.

In this study, we present our work on Taxor and show benchmarking results of a comparison with five commonly used metagenomic profiling tools. Taxor’s read classification results are on par with state-of-the-art methods regarding recall, while improving precision rates in almost every experiment on real data. Our new tool consistently shows the highest *F*_1_and *F*_0.5_-scores on the species level for the three mock communities, indicating the robustness of the findings. We attribute this improvement to using syncmers instead of minimizers, applying several iterative filter steps, and our EM algorithm for read classification refinement. This assumption is based on the observation that Taxor and KMCP perform best regarding precision. Both use syncmers, similar filter steps, and a final EM algorithm for the profiling. They mainly differ in the underlying data structure for approximate membership querying and their implementation of the EM algorithm.

Contamination and environmental DNA are important aspects in many metagenomic studies, particularly if their genomes are not in the databases used by the metagenomic profilers. We have shown that Taxor is robust against these out-of-database genomes, minimizing the risk of false classifications of reads in such scenarios. Here, Taxor outperforms the mapping-based method MetaMaps and is on par with the Bloom Filter approaches of Ganon and KMCP. The false positive rate of 0.3%, which we used in this study for the Bloom Filter approaches, and our HIXF data structure seems to protect the three tools from falsely classifying reads from out-of-database genomes. Our observations suggest that removing host reads may not be necessary for these tools before the metagenomic classification of long reads in microbiome studies.

Correctly estimating the composition of microbial genomes in metagenomic samples is a main task in many microbiome studies, investigating the differential abundances of species between several gut or environmental samples. These differences can be attributed to environmental changes like global warming or anthropogenic pollution. Our benchmarking results show that none of the evaluated tools accurately estimates the abundance of all species in the real mock communities used for evaluation. Although Taxor also has problems estimating the species abundances correctly, its calculated sequence and taxonomic abundances are more consistent with the theoretical abundances of species in the communities. However, these results should be taken with a pinch of salt as various factors, including different DNA extraction methods, can affect the final composition of DNA sequenced for metagenomic samples and potentially bias relative abundance estimates of the tools (49).

The small MinION sequencing devices invented by Oxford Nanopore Technologies (ONT) provide the possibility to sequence a sample at the place of its origin without the need to ship the sample to a laboratory. In such a point-of-care sequencing scenario, computing resources for metagenomic analysis are usually limited. Without a reliable internet connection to perform analysis in the cloud, the tools applied for metagenomic profiling must be as fast and memory efficient as possible while retaining high read classification accuracy. In this context, Taxor represents a significant improvement over state-of-the-art tools, requiring only 50% of the memory and disk space as the best competitor in our benchmarking. Taxor was also among the fastest tools regarding the query time while showing the highest average precision across the three real evaluation datasets. We expect Taxor to be a valuable tool for usage in real-time long-read metagenomic analysis pipelines like minoTour (50) or WIMP (42), both using Centrifuge for read classification.

Our new HIXF data structure significantly improves the interleaved Bloom filter concerning memory consumption and query time. It also reduces the memory requirements compared to the recently published hierarchical interleaved Bloom filter (29) when both use the same false positive rate. However, this comes at the cost of less flexibility and more time needed to build the index. The biggest drawback of the XOR filter is the ability to only index static sets of keys. That means all input data need to be known a priori, and the index cannot be updated after it has been built once. Since the XOR filter is the underlying data structure of our HIXF, this also applies to it. However, we argue that build times for the HIXF are still acceptable and that the lower memory footprint for querying reads compensates for that issue. We even envisage that tools currently relying on Bloom filters or interleaved Bloom filters can benefit from utilizing the HIXF as their index data structure, particularly in time-critical applications like nanopore adaptive sampling (51).

There are two important directions for the future development of Taxor. First, reducing the computational requirements by enhancing the underlying data structure is worth the effort because the number of genomes in public databases is constantly growing. One possibility would be using Binary Fuse or Ribbon filters instead of the XOR Filter in the hierarchical interleave data structure. Recent studies have shown that both filter types are practically smaller than XOR filters (33, 52). Secondly, estimating the species abundances in the investigated metagenomic samples needs to be improved to reliably use our tool in microbiome studies relying on differential abundance calculations. Here, enhancing the profiling would require further filtering and an improvement of our EM algorithm, which remains an open task for further research studies.

## Conclusions

This study presents Taxor, a long-read metagenomic read classification and profiling tool that uses hierarchical interleaved XOR filters for indexing and querying large genome databases. Taxor is competitive with state-of-the-art metagenomic classifiers regarding precision and recall with the advantage of being more memory-efficient. When querying hundreds of thousands of long reads against the index, Taxor is on par with the fastest competitors reducing memory and disk space by more than 50%.

## Availability of data and materials

We have made the source code of Taxor available as open-source software at https://gitlab.com/dacs-hpi/taxor. Output files of the compared tools, and Python and R Scripts used for data analysis are available at https://osf.io/vqkcn/.

## Funding

This work was funded by the German Federal Ministry of Education and Research (BMBF) in the Computational Life Science program (Live-DREAM, 031L0175B) and Global Health program (ZooSeq, 01KI1905D) and has been supported by a grant from the BMBF/German Center for Infection Research (TI 06.904 - FP2019 to BYR).

## ACKNOWLEDGEMENTS

The authors thank Vitor Piro, Enrico Seiler, and Knut Reinert (FU Berlin) for valuable discussions and comments on hierarchical filters and metagenomic profiling.

## Notes

https://gitlab.com/dacs-hpi/taxor

https://osf.io/vqkcn/

### Competing Interest Statement

The authors have declared no competing interest.

## Bibliography

1. Andreas Andrusch, Piotr W Dabrowski, Jeanette Klenner, Simon H Tausch, Claudia Kohl, Abdalla A Osman, Bernhard Y Renard, and Andreas Nitsche. Paipline: pathogen identification in metagenomic and clinical next generation sequencing samples. Bioinformatics, 34 (17):i715–i721, 2018.

2. Vesselin V Doytchinov and Svetoslav G Dimov. Microbial community composition of the antarctic ecosystems: Review of the bacteria, fungi, and archaea identified through an ngs-based metagenomics approach. Life, 12(6):916, 2022.

3. Daehwan Kim, Li Song, Florian P Breitwieser, and Steven L Salzberg. Centrifuge: rapid and sensitive classification of metagenomic sequences. Genome research, 26(12):1721–1729, 2016.

4. Derrick E Wood, Jennifer Lu, and Ben Langmead. Improved metagenomic analysis with kraken 2. Genome biology, 20:1–13, 2019.

5. Alexander T Dilthey, Chirag Jain, Sergey Koren, and Adam M Phillippy. Strain-level metagenomic assignment and compositional estimation for long reads with metamaps. Nature communications, 10(1):3066, 2019.

6. Martin S Lindner and Bernhard Y Renard. Metagenomic abundance estimation and diagnostic testing on species level. Nucleic acids research, 41(1):e10–e10, 2013.

7. Martina Fischer, Benjamin Strauch, and Bernhard Y Renard. Abundance estimation and differential testing on strain level in metagenomics data. Bioinformatics, 33(14):i124–i132, 2017.

8. Temesgen Hailemariam Dadi, Bernhard Y Renard, Lothar H Wieler, Torsten Semmler, and Knut Reinert. Slimm: species level identification of microorganisms from metagenomes. PeerJ, 5:e3138, 2017.

9. Vitor C Piro, Martin S Lindner, and Bernhard Y Renard. Dudes: a top-down taxonomic profiler for metagenomics. Bioinformatics, 32(15):2272–2280, 2016.

10. Changjin Hong, Solaiappan Manimaran, Ying Shen, Joseph F Perez-Rogers, Allyson L Byrd, Eduardo Castro-Nallar, Keith A Crandall, and William Evan Johnson. Pathoscope 2.0: a complete computational framework for strain identification in environmental or clinical sequencing samples. Microbiome, 2(1):1–15, 2014.

11. Ben Langmead and Steven L Salzberg. Fast gapped-read alignment with bowtie 2. Nature methods, 9(4):357–359, 2012.

12. Duy Tin Truong, Eric A Franzosa, Timothy L Tickle, Matthias Scholz, George Weingart, Edoardo Pasolli, Adrian Tett, Curtis Huttenhower, and Nicola Segata. Metaphlan2 for enhanced metagenomic taxonomic profiling. Nature methods, 12(10):902–903, 2015.

13. Alessio Milanese, Daniel R Mende, Lucas Paoli, Guillem Salazar, Hans-Joachim Ruscheweyh, Miguelangel Cuenca, Pascal Hingamp, Renato Alves, Paul I Costea, Luis Pedro Coelho, et al. Microbial abundance, activity and population genomic profiling with motus2. Nature communications, 10(1):1014, 2019.

14. Qiaoxing Liang, Paul W Bible, Yu Liu, Bin Zou, and Lai Wei. Deepmicrobes: taxonomic classification for metagenomics with deep learning. NAR Genomics and Bioinformatics, 2 (1):lqaa009, 2020.

15. Florian Mock, Fleming Kretschmer, Anton Kriese, Sebastian Böcker, and Manja Marz. Taxonomic classification of dna sequences beyond sequence similarity using deep neural networks. Proceedings of the National Academy of Sciences, 119(35):e2122636119, 2022.

16. Vitor C Piro, Temesgen H Dadi, Enrico Seiler, Knut Reinert, and Bernhard Y Renard. ganon: precise metagenomics classification against large and up-to-date sets of reference sequences. Bioinformatics, 36(Supplement_1):i12–i20, 2020.

17. Wei Shen, Hongyan Xiang, Tianquan Huang, Hui Tang, Mingli Peng, Dachuan Cai, Peng Hu, and Hong Ren. Kmcp: accurate metagenomic profiling of both prokaryotic and viral populations by pseudo-mapping. Bioinformatics, 39(1):btac845, 2023.

18. Nuala A O’Leary, Mathew W Wright, J Rodney Brister, Stacy Ciufo, Diana Haddad, Rich McVeigh, Bhanu Rajput, Barbara Robbertse, Brian Smith-White, Danso Ako-Adjei, et al. Reference sequence (refseq) database at ncbi: current status, taxonomic expansion, and functional annotation. Nucleic acids research, 44(D1):D733–D745, 2016.

19. Donovan H Parks, Maria Chuvochina, Christian Rinke, Aaron J Mussig, Pierre-Alain Chaumeil, and Philip Hugenholtz. Gtdb: an ongoing census of bacterial and archaeal diversity through a phylogenetically consistent, rank normalized and complete genome-based taxonomy. Nucleic acids research, 50(D1):D785–D794, 2022.

20. Guillaume Holley and Páll Melsted. Bifrost: highly parallel construction and indexing of colored and compacted de bruijn graphs. Genome biology, 21(1):1–20, 2020.

21. Prashant Pandey, Fatemeh Almodaresi, Michael A Bender, Michael Ferdman, Rob Johnson, and Rob Patro. Mantis: a fast, small, and exact large-scale sequence-search index. Cell systems, 7(2):201–207, 2018.

22. Burton H Bloom. Space/time trade-offs in hash coding with allowable errors. Communications of the ACM, 13(7):422–426, 1970.

23. Robert S Harris and Paul Medvedev. Improved representation of sequence bloom trees. Bioinformatics, 36(3):721–727, 2020.

24. Brad Solomon and Carl Kingsford. Improved search of large transcriptomic sequencing databases using split sequence bloom trees. Journal of Computational Biology, 25(7):755–765, 2018.

25. Chen Sun, Robert S Harris, Rayan Chikhi, and Paul Medvedev. Allsome sequence bloom trees. Journal of Computational Biology, 25(5):467–479, 2018.

26. Phelim Bradley, Henk C Den Bakker, Eduardo PC Rocha, Gil McVean, and Zamin Iqbal. Ultrafast search of all deposited bacterial and viral genomic data. Nature biotechnology, 37 (2):152–159, 2019.

27. Timo Bingmann, Phelim Bradley, Florian Gauger, and Zamin Iqbal. Cobs: a compact bitsliced signature index. In String Processing and Information Retrieval: 26th International Symposium, SPIRE 2019, Segovia, Spain, October 7–9, 2019, Proceedings 26, pages 285–303. Springer, 2019.

28. Temesgen Hailemariam Dadi, Enrico Siragusa, Vitor C Piro, Andreas Andrusch, Enrico Seiler, Bernhard Y Renard, and Knut Reinert. Dream-yara: An exact read mapper for very large databases with short update time. Bioinformatics, 34(17):i766–i772, 2018.

29. Svenja Mehringer, Enrico Seiler, Felix Droop, Mitra Darvish, René Rahn, Martin Vingron, and Knut Reinert. Hierarchical interleaved bloom filter: Enabling ultrafast, approximate sequence queries. bioRxiv, pages 2022–08, 2022.

30. Thomas Mueller Graf and Daniel Lemire. Xor filters: Faster and smaller than bloom and cuckoo filters. Journal of Experimental Algorithmics (JEA), 25:1–16, 2020.

31. Bin Fan, Dave G Andersen, Michael Kaminsky, and Michael D Mitzenmacher. Cuckoo filter: Practically better than bloom. In Proceedings of the 10th ACM International on Conference on emerging Networking Experiments and Technologies, pages 75–88, 2014.

32. Michael Mitzenmacher, Salvatore Pontarelli, and Pedro Reviriego. Adaptive cuckoo filters, 2020.

33. Thomas Mueller Graf and Daniel Lemire. Binary fuse filters: Fast and smaller than xor filters. Journal of Experimental Algorithmics (JEA), 27(1):1–15, 2022.

34. Knut Reinert, Temesgen Hailemariam Dadi, Marcel Ehrhardt, Hannes Hauswedell, Svenja Mehringer, René Rahn, Jongkyu Kim, Christopher Pockrandt, Jörg Winkler, Enrico Siragusa, et al. The seqan c++ template library for efficient sequence analysis: A resource for programmers. Journal of biotechnology, 261:157–168, 2017.

35. Robert Edgar. Syncmers are more sensitive than minimizers for selecting conserved k-mers in biological sequences. PeerJ, 9:e10805, 2021.

36. Abhinav Dutta, David Pellow, and Ron Shamir. Parameterized syncmer schemes improve long-read mapping. PLOS Computational Biology, 18(10):e1010638, 2022.

37. Fabiano C Botelho, Rasmus Pagh, and Nivio Ziviani. Simple and space-efficient minimal perfect hash functions. In Algorithms and Data Structures: 10th International Workshop, WADS 2007, Halifax, Canada, August 15-17, 2007. Proceedings 10, pages 139–150. Springer, 2007.

38. Philippe Flajolet, Éric Fusy, Olivier Gandouet, and Frédéric Meunier. Hyperloglog: the analysis of a near-optimal cardinality estimation algorithm. In Discrete Mathematics and Theoretical Computer Science, pages 137–156. Discrete Mathematics and Theoretical Computer Science, 2007.

39. Daniel N Baker and Ben Langmead. Dashing: fast and accurate genomic distances with hyperloglog. Genome biology, 20:1–12, 2019.

40. Ryan R Wick. Badread: simulation of error-prone long reads. Journal of Open Source Software, 4(36):1316, 2019.

41. Chi-Ming Leung, Dinghua Li, Yan Xin, Wai-Chun Law, Yifan Zhang, Hing-Fung Ting, Ruibang Luo, and Tak-Wah Lam. Megapath: sensitive and rapid pathogen detection using metagenomic ngs data. BMC genomics, 21(6):1–9, 2020.

42. Sissel Juul, Fernando Izquierdo, Adam Hurst, Xiaoguang Dai, Amber Wright, Eugene Kulesha, Roger Pettett, and Daniel J Turner. What’s in my pot? real-time species identification on the minion™. BioRxiv, page 030742, 2015.

43. Yukiteru Ono, Kiyoshi Asai, and Michiaki Hamada. Pbsim2: a simulator for long-read sequencers with a novel generative model of quality scores. Bioinformatics, 37(5):589–595, 2021.

44. Scott Ferguson, Todd McLay, Rose L Andrew, Jeremy J Bruhl, Benjamin Schwessinger, Justin Borevitz, and Ashley Jones. Species-specific basecallers improve actual accuracy of nanopore sequencing in plants. Plant Methods, 18(1):1–11, 2022.

45. Samuel M Nicholls, Joshua C Quick, Shuiquan Tang, and Nicholas J Loman. Ultra-deep, long-read nanopore sequencing of mock microbial community standards. Gigascience, 8 (5):giz043, 2019.

46. Heng Li. Minimap2: pairwise alignment for nucleotide sequences. Bioinformatics, 34(18): 3094–3100, 2018.

47. Zheng Sun, Shi Huang, Meng Zhang, Qiyun Zhu, Niina Haiminen, Anna Paola Carrieri, Yoshiki Vázquez-Baeza, Laxmi Parida, Ho-Cheol Kim, Rob Knight, et al. Challenges in benchmarking metagenomic profilers. Nature methods, 18(6):618–626, 2021.

48. Daniel M Portik, C Titus Brown, and N Tessa Pierce-Ward. Evaluation of taxonomic classification and profiling methods for long-read shotgun metagenomic sequencing datasets. BMC bioinformatics, 23(1):541, 2022.

49. Hui-yu Sui, Ana A Weil, Edwin Nuwagira, Firdausi Qadri, Edward T Ryan, Melissa P Mezzari, Wanda Phipatanakul, and Peggy S Lai. Impact of dna extraction method on variation in human and built environment microbial community and functional profiles assessed by shotgun metagenomics sequencing. Frontiers in microbiology, 11:953, 2020.

50. Rory Munro, Roberto Santos, Alexander Payne, Teri Forey, Solomon Osei, Nadine Holmes, and Matthew Loose. minotour, real-time monitoring and analysis for nanopore sequencers. Bioinformatics, 38(4):1133–1135, 2022.

51. Jens-Uwe Ulrich, Ahmad Lutfi, Kilian Rutzen, and Bernhard Y Renard. ReadBouncer: precise and scalable adaptive sampling for nanopore sequencing. Bioinformatics, 38 (Supplement_1):i153.#x2013;i160, 06 2022.

52. Peter C Dillinger and Stefan Walzer. Ribbon filter: practically smaller than bloom and xor. arXiv preprint arXiv:2103.02515, 2021.

