## Supplemental Material for "Taxor: Fast and space-efficient taxonomic classification of long reads with hierarchical interleaved XOR filters"

**for**

### Threshold calculation for the k-mer based model

For our new HIXF data structure, we use membership queries for all k-mers of a given read against all k-mers of a given reference sequence set. We consider a read a hit with a reference sequence if the number of matching k-mers is greater than or equal to a given threshold $t$. For the k-mer model, we calculate the threshold using the expected sequencing error rate e and the definition of a $(1-\alpha)$ confidence interval of the number of erroneous k-mers as provided by Blanca et al. For a given read $r$ with length $len(r)$ and k-mer length $k$, we denote the number of k-mers of a read $r$ as $L = len(r) - k + 1$, and define $q$ by $(1-(1-e)^k)$. Then, the expected number of erroneous k-mers can be calculated as follows:

$$E[N_{err}] = L \times q$$

Let $Var(N_{err})$ be the variance for the number of erroneous k-mers. We can then calculate the upper bound of the $(1-\alpha)$ confidence interval by

$$E[N_{err}] + z_\alpha \sqrt{Var(N_{err})}$$

With $z_\alpha = \phi^{-1}(1-\frac{\alpha}{2})$, where we denote $\phi^{-1}$ as the inverse of the cumulative distribution function of the standard Gaussian distribution. Based on the calculation of the confidence interval for the number of erroneous k-mers, we define our threshold for the minimum number of matching k-mers for read $r$ as:

$$\min[N_{match}] = L - (E[N_{err}] + z_\alpha \sqrt{Var(N_{err})})$$

We classify a read as a match with a reference sequence if the number of matching k-mers is bigger or equal to

$$t = \min[N_{match}]$$

### Profiling methods commands

To facilitate reproducible results, we provide the general commands or instructions to run each method.

#### genome_updater

We used genome_updater version 0.5.2 to download all complete genome sequences and chromosomes of archaea, bacteria, viruses, and fungi from the NCBI RefSeq database (Release 217)

```
genome_updater.sh –d refseq –g archaea,bacteria,fungi,viral -l complete genome,chromosome, -f genomic.fna.gz -o refseq-abfv –t 12 -A species:1 -m –a –p

mkdir –p refseq-abfv/2023-03-15_12-56-12/taxdump

tar –xvzf refseq-abfv/2023-03-15_12-56-12/taxdump.tar.gz –C refseq-abfv/2023-03-15_12-56-12/taxdump
```

#### Centrifuge

We ran Centrifuge version 1.0.4. First we created a Fasta file of all downloaded reference genomes and a corresponding conversion table. Then we can use the prepared input data and the downloaded NCBI taxonomy dump to build a Centrifuge database index

```
cut –f 1,6 refseq-abfv/2023-03-15_12-56-12/assembly_summary.txt > refseq-abfv/2023-03-15_12-56-12/taxid.map

centrifuge_conversion_table.py -t refseq-abfv/2023-03-15_12-56-12/taxid.map -s refseq-abfv/2023-03-15_12-56-12/files -o refseq-abfv/2023-03-15_12-56-12/centrifuge_data/conversion_table.tsv

centrifuge-build --conversion-table refseq-abfv/2023-03-15_12-56-12/centrifuge_data/conversion_table.tsv --taxonomy-tree refseq-abfv/2023-03-15_12-56-12/taxdump/nodes.dmp --name-table refseq-abfv/2023-03-15_12-56-12/taxdump/names.dmp -p 30 refseq-abfv/2023-03-15_12-56-12/files/all.fna refseq-abfv/2023-03-15_12-56-12/centrifuge_data/refseq-abfv
```

Finally, we query the fastq file of one of the samples against the index and create a kraken report file from the Centrifuge output.

```
centrifuge -q --min-hitlen 22 -k 20 -t -p 30 -x refseq-abfv/2023-03-15_12-56-12/centrifuge_data/refseq-abfv -U SAMPLE.fastq.gz -S refseq-abfv/2023-03-15_12-56-12/centrifuge_data/SAMPLE.centrifuge.search.txt --report-file refseq-abfv/2023-03-15_12-56-12/centrifuge_data/SAMPLE.centrifuge.report.tsv

centrifuge-kreport -x refseq-abfv/2023-03-15_12-56-12/centrifuge_data/refseq-abfv --no-lca refseq-abfv/2023-03-15_12-56-12/centrifuge_data/SAMPLE.centrifuge.search.txt > refseq-abfv/2023-03-15_12-56-12/centrifuge_data/SAMPLE.centrifuge.kreport.txt
```

### Kraken2

We ran Kraken2 version 2.1.2. We first needed to prepare input data by adding the Kraken header information to the fasta file using the conversion table we also created for building the Centrifuge index.

```
kraken2-build --download-taxonomy --db  refseq-abfv/2023-03-15_12-56-12/kraken2_data/refseq-abfv

add_kraken_header.py -t refseq-abfv/2023-03-15_12-56-12/centrifuge_data/conversion_table.tsv -f refseq-abfv/2023-03-15_12-56-12/files/all.fna -o refseq-abfv/2023-03-15_12-56-12/kraken2_data/all_seq.fna

kraken2-build -add-to-library refseq-abfv/2023-03-15_12-56-12/kraken2_data/all_seq.fna --db refseq-abfv/2023-03-15_12-56-12/kraken2_data/refseq-abfv-k32-m22 --no-masking

kraken2-build --build --kmer-len 32 --minimizer-len 22 --minimizer-spaces 0 --threads 30 --db refseq-abfv/2023-03-15_12-56-12/kraken2_data/refseq-abfv-k32-m22
```

After building the database index we can query each sample against it, resulting in a kraken report file and an output txt with binning information per read.

```
kraken2 --db refseq-abfv/2023-03-15_12-56-12/kraken2_data/refseq-abfv-k32-m22 --threads 30
--report refseq-abfv/2023-03-15_12-56-12/kraken2_data/SAMPLE.report --output refseq-
abfv/2023-03-15_12-56-12/kraken2_data/SAMPLE.output.txt --gzip-compressed SAMPLE.fq.gz
```

### KMCP

We used KMCP version 0.9.1. First, we needed to prepare the input data for creating the database index, as described in the KMCP wiki (https://bioinf.shenwei.me/kmcp/database/#refseq-viral-or-fungi) using the tools `rush` and `brename`, provided by the author of KMCP

```
cut -f 1,6 refseq-abfv/2023-03-15_12-56-12/assembly_summary.txt > refseq-abfv/2023-03-
15_12-56-12/taxid.map

cut -f 1,8 refseq-abfv/2023-03-15_12-56-12/assembly_summary.txt > refseq-abfv/2023-03-
15_12-56-12/name.map

mkdir -p refseq-abfv/2023-03-15_12-56-12/kmcp_data/files.renamed

cd refseq-abfv/2023-03-15_12-56-12/kmcp_data/files.renamed

find refseq-abfv/2023-03-15_12-56-12/files -name "*.fna.gz" | rush 'ln -s {}'

cd ..

brename -R -p '^(\w{3}_\d{9}\.\d+).+' -r '$1.fna.gz' refseq-abfv/2023-03-15_12-56-
12/kmcp_data/files.renamed
```

In the next step, we compute the syncmers and create the database index with a false positive rate of 0.3%.

```
kmcp compute -I refseq-abfv/2023-03-15_12-56-12/kmcp_data/files.renamed -O refseq-
abfv/2023-03-15_12-56-12/kmcp_data/refseq-abfv-k22-s12 -S 12 -k 22 --seq-name-filter
plasmid --split-number 10 --split-overlap 150 --log refseq-abfv-k22-s12.log -j 30 --force

kmcp index -I refseq-abfv/2023-03-15_12-56-12/kmcp_data/refseq-abfv-k22-s12/ -O refseq-
abfv/2023-03-15_12-56-12/kmcp_data/refseq-abfv-k22-s12.kmcp -j 30 -f 0.003 -n 3 -x 100K --
log refseq-abfv-k22-s12.kmcp.log --force
```

Finally, the sample file is queried against the index and the profiling refines read assignments and report taxonomic abundances.

```
kmcp search --db-dir refseq-abfv/2023-03-15_12-56-12/kmcp_data/refseq-abfv-k22-s12.kmcp
--threads 30 -f 0.003 --min-query-cov 0.12 --out-file refseq-abfv/2023-03-15_12-56-
12/kmcp_data/SAMPLE.tsv.gz SAMPLE.fq.gz

kmcp profile --taxid-map refseq-abfv/2023-03-15_12-56-12/kmcp_data/taxid.map --taxdump
refseq-abfv/2023-03-15_12-56-12/taxdump/ --level species --min-query-cov 0.12 -m 3 refseq-
abfv/2023-03-15_12-56-12/kmcp_data/SAMPLE.tsv.gz --min-hic-ureads-qcov 0.2 --min-chunks-
fraction 0.2 --out-prefix refseq-abfv/2023-03-15_12-56-12/kmcp_data/SAMPLE.kmcp.profile
```

```
--cami-report refseq-abfv/2023-03-15_12-56-12/kmcp_data/SAMPLE.cami.profile --sample-id
SAMPLE_NAME --binning-result refseq-abfv/2023-03-15_12-56-12/kmcp_data/SAMPLE.binning.gz
```

### Ganon

We used Ganon version 1.5.0. Here, no further preprocessing step is needed to create the custom database index. We used the same minimizer and k-mer lengths as for Kraken2 and created the index with a false positive rate of 0.3%. When classifying the SAMPLE reads, ganon reports read assignments and taxonomic abundances in CAMI report format.

```
ganon build-custom --db-prefix refseq-abfv/2023-03-15_12-56-12/ganon_data/refseq-abfv
--input refseq-abfv/2023-03-15_12-56-12/files/ --level species --ncbi-file-info refseq-
abfv/2023-03-15_12-56-12/assembly_summary.txt --threads 30 --max-fp 0.003 --kmer-size 22
--window-size 32 --hash-functions 3

ganon classify --db-prefix refseq-abfv/2023-03-15_12-56-12/ganon_data/refseq-abfv
-s SAMPLE.fq.gz -o refseq-abfv/2023-03-15_12-56-12/ganon_data/SAMPLE.search --threads 30
-a --output-all -c 0.12 -e 0.9
```

### MetaMaps

We used MetaMaps version 0.1. To create the custom database index, we followed the steps described at https://github.com/DiltheyLab/MetaMaps#databases.

```
mkdir refseq-abfv/2023-03-15_12-56-12/metamaps_data/download

perl downloadRefSeq.pl --seqencesOutDirectory refseq-abfv/2023-03-15_12-56-
12/metamaps_data/download/refseq --taxonomyOutDirectory refseq-abfv/2023-03-15_12-56-
12/metamaps_data/download/taxonomy –targetBranches archae,bacteria,fungi,viral
```

Then, we needed to modify the following line in script `annotateRefSeqSequencesWithUniqueTaxonIDs.pl` from

```
next unless($assembly_level eq 'Complete Genome');
```
to
```
next unless($assembly_level eq 'Complete Genome' || ($assembly_level eq 'Chromosome');
```

and execute the script using the taxonomy downloaded by genome_updater and build the database used for indexing.

```
perl annotateRefSeqSequencesWithUniqueTaxonIDs.pl --refSeqDirectory refseq-abfv/2023-03-
15_12-56-12/metamaps_data/download/refseq --taxonomyInDirectory refseq-abfv/2023-03-15_12-
56-12/taxdump/ --taxonomyOutDirectory refseq-abfv/2023-03-15_12-56-
12/metamaps_data/download/taxonomy_uniqueIDs

mkdir –p refseq-abfv/2023-03-15_12-56-12/taxdump/ --taxonomyOutDirectory refseq-abfv/2023-
03-15_12-56-12/metamaps_data/databases
```

```
perl buildDB.pl --DB refseq-abfv/2023-03-15_12-56-12/metamaps_data/databases/refseq-abfv
--FASTAs refseq-abfv/2023-03-15_12-56-12/metamaps_data/download/refseq --taxonomy refseq-
abfv/2023-03-15_12-56-12/metamaps_data/download/taxonomy_uniqueIDs
```

Then we finally index the created database with the following command.

```
metamaps index -r refseq-abfv/2023-03-15_12-56-12/metamaps_data/databases/refseq-
abfv/DB.fa -t 30 -i refseq-abfv/2023-03-15_12-56-12/metamaps_data/refseq-abfv-k16
```

For querying sample reads against the created index, we used the following commands

```
metamaps mapAgainstIndex --all -q SAMPLE.fq.gz -i refseq-abfv/2023-03-15_12-56-
12/metamaps_data/databases/refseq-abfv-k16 -o refseq-abfv/2023-03-15_12-56-
12/metamaps_data/SAMPLE.map.txt -t 30

metamaps classify --DB refseq-abfv/2023-03-15_12-56-12/metamaps_data/databases/refseq-
abfv-k16 -t 30 --mappings refseq-abfv/2023-03-15_12-56-12/metamaps_data/SAMPLE.map.txt
```

### Taxor

We used Taxor version 0.1.0. For preprocessing, we need to create a tab-separated file containing important taxonomic information, using the tool taxonkit (https://github.com/shenwei356/taxonkit).

```
cut -f 1,7,20 refseq-abfv/2023-03-15_12-56-12/assembly_summary.txt | taxonkit lineage -i 2
-r -n -L --data-dir refseq-abfv/2023-03-15_12-56-12/taxdump | taxonkit reformat -I 2 -P -t
--data-dir refseq-abfv/2023-03-15_12-56-12/taxdump | cut -f 1,2,3,4,6,7 > refseq-
abfv/2023-03-15_12-56-12/taxor_data/refseq_accessions_taxonomy.csv
```

Then we build the Taxor index using the tab separated file and the sequence files downloaded with genome_updater.

```
taxor build --input-file refseq-abfv/2023-03-15_12-56-
12/taxor_data/refseq_accessions_taxonomy.csv --input-sequence_dir refseq-abfv/2023-03-
15_12-56-12/files --output-filename refseq-abfv/2023-03-15_12-56-12/taxor_data/refseq-
abfv-k22-s12.hixf --threads 30 --kmer-size 22 --syncmer-size 12 --use-syncmer
```

Finally, we query the sample fastq file against the index allowing a sequencing error rate of 15%. The query result file is used as input for taxonomic profiling, which has three output files containing taxonomic abundances and sequence abundances in CAMI report format as well as a binning file with final read to reference assignments.

```
taxor search --index-file refseq-abfv/2023-03-15_12-56-12/taxor_data/refseq-abfv-k22-
s12.hixf --query-file SAMPLE.fq.gz --output-file refseq-abfv/2023-03-15_12-56-
12/taxor_data/SAMPLE.search.txt --error-rate 0.15 --threads 30

taxor profile --search-file refseq-abfv/2023-03-15_12-56-12/taxor_data/SAMPLE.search.txt
--cami-report-file refseq-abfv/2023-03-15_12-56-12/taxor_data/SAMPLE.report
--seq-abundance-file refseq-abfv/2023-03-15_12-56-12/taxor_data/SAMPLE.abundance
--binning-file refseq-abfv/2023-03-15_12-56-12/taxor_data/SAMPLE.binning --sample-id
SAMPLE
```
